# Phytoplankton thermal responses adapt in the absence of hard thermodynamic constraints

**DOI:** 10.1101/452250

**Authors:** Dimitrios - Georgios Kontopoulos, Erik van Sebille, Michael Lange, Gabriel Yvon-Durocher, Timothy G. Barraclough, Samraat Pawar

## Abstract

To better predict how populations and communities respond to climatic temperature variation, it is necessary to understand how the shape of the response of fitness-related traits to temperature evolves (the thermal performance curve). Currently, there is disagreement about the extent to which the evolution of thermal performance curves is constrained. One school of thought has argued for the prevalence of thermodynamic constraints through enzyme kinetics, whereas another argues that adaptation can—at least partly—overcome such constraints. To shed further light on this debate, we perform a phylogenetic meta-analysis of the thermal performance curves of growth rate of phytoplankton—a globally important functional group—, controlling for environmental effects (habitat type and thermal regime). We find that thermodynamic constraints have a minor influence on the shape of the curve. In particular, we detect a very weak increase of maximum performance with the temperature at which the curve peaks, suggesting a weak “hotter-is-better” constraint. Also, instead of a constant thermal sensitivity of growth across species, as might be expected from strong constraints, we find that all aspects of the thermal performance curve evolve along the phylogeny. Our results suggest that phytoplankton thermal performance curves adapt to thermal environments largely in the absence of hard thermodynamic constraints.

## Introduction

Temperature changes can affect the dynamics of all levels of biological organization by changing the metabolic rate of individual organisms (Brown et al., 2004; Pörtner et al., 2006; Hoffmann and Sgró, 2011; Pawar et al., 2015). Thus, to better understand the impacts of current and future climate change on whole ecosystems, it is essential to understand how key fitness-related metabolic traits (e.g., growth rate, photosynthesis rate) respond to changes in environmental temperature.

In ectotherms, the relationship of fitness-related traits with temperature (the “thermal performance curve”; TPC) is typically unimodal (Fig. 1; Angilletta 2009). Trait values increase with temperature until a critical point (*T*_pk_), after which they drop rapidly. To understand the capacity for adaptation of the TPC to different thermal environments, it is important to investigate how the shape of the TPC evolves across species and, in so doing, to identify any constraining factors that operate over both short (ecological and microevolutionary) and long (macroevolutionary) timescales. This remains an area of ongoing debate, with multiple competing hypotheses existing in the literature. Such hypotheses can be broadly classified along a continuum that ranges from strong and insurmountable constraints on TPC evolution due to thermodynamic constraints on enzyme kinetics, to weak constraints that can be overcome through adaptation (Fig. 2).

**Figure 1.**
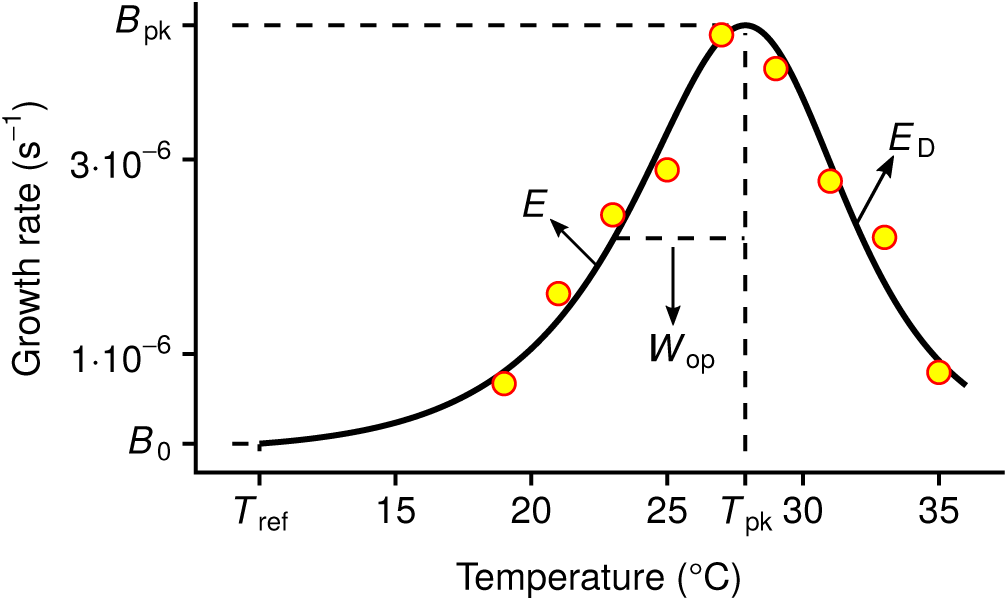
The relationship of growth rate (*r*_max_) with temperature in ectotherms (the thermal performance curve; TPC). The TPC is generally unimodal and asymmetric, here quantified by the four-parameter Sharpe-Schoolfield model (black line; Schoolfield et al. 1981) fitted to growth rate measurements of the dinoflagellate *Amphidinium klebsii* (Morton et al., 1992). The parameters of the model are *B*_0_ (in units of s^−1^), *E* (eV), *T*_pk_ (K), and *E*_D_ (eV). *B*_0_ is the growth rate at a reference temperature below the peak (*T*_ref_) and controls the vertical offset of the TPC. *E* sets the rate at which the curve rises and is, therefore, a measure of thermal sensitivity at the operational temperature range. *T*_pk_ is the temperature at which growth rate is maximal, and *E*_D_ controls the fall of the curve. Two other parameters control the shape of the curve and can be calculated from the four main parameters: *B*_pk_ (s^−1^); the maximum height of the curve, and *W*_op_ (K); the operational niche width, which we define as the difference between *T*_pk_ and the temperature at the rise of the curve where growth rate is half of *B*_pk_. Further information regarding the assumptions of the model are provided in the section “Estimation of TPC parameter values” of the Methods.

**Figure 2.**
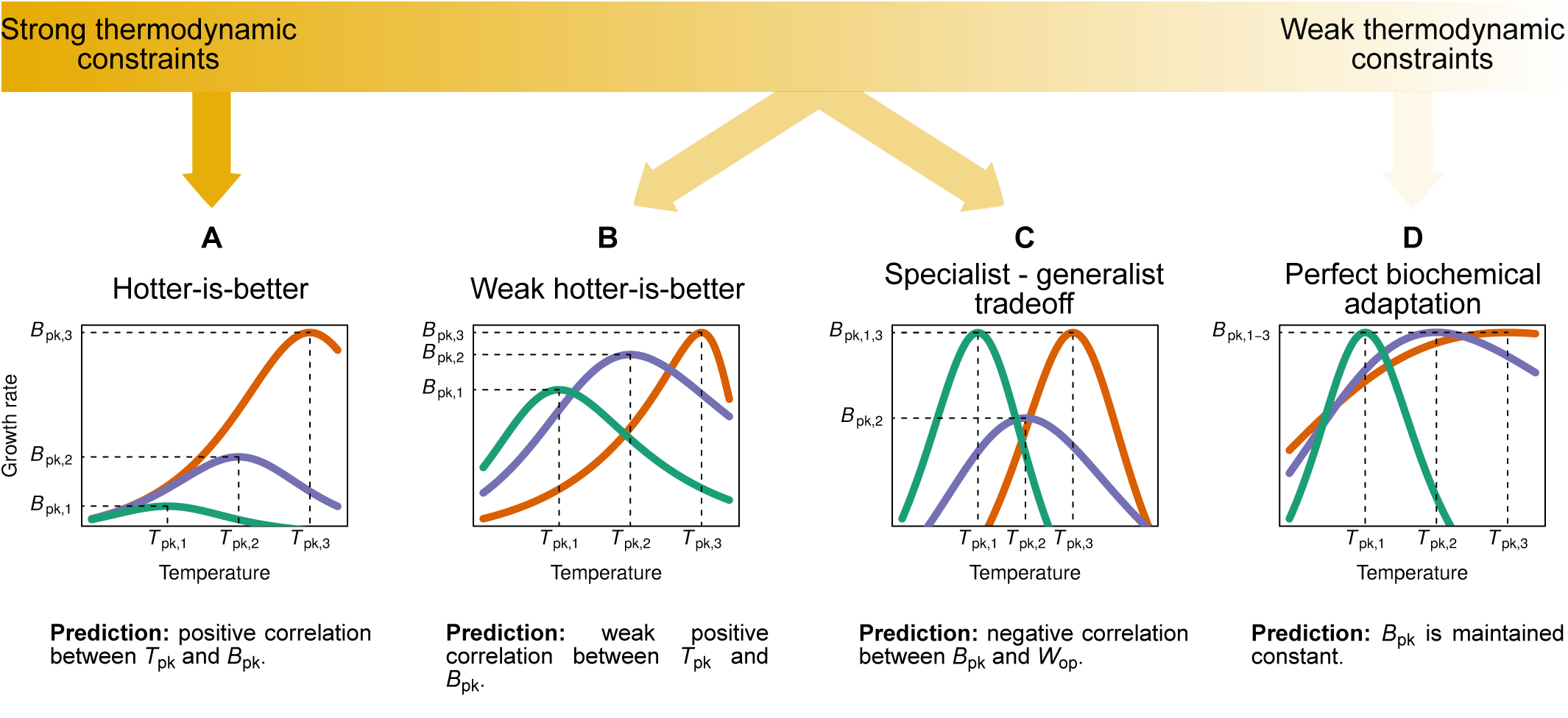
The spectrum of hypotheses for the evolution of thermal performance curves across species. A key area of difference among these hypotheses concerns the impact of thermodynamic constraints on the shape of the TPC. Thus, hypotheses can be classified between those lying near the strong thermodynamic constraints end, the middle of the spectrum, or the weak thermodynamic constraints end where thermodynamic constraints can be overcome through biochemical adaptation. It is worth clarifying that in panel D, the maximum value that *B*_pk_ can take would also be under a thermodynamic constraint, but this constraint would be different from those assumed in panels A and B. A detailed description of each hypothesis and its assumptions (e.g., the impact of body size on trait performance) is provided in the main text.

At the strong thermodynamic constraints extreme, the “hotter-is-better” hypothesis (Frazier et al., 2006; Kingsolver and Huey, 2008; Knies et al., 2009; Angilletta et al., 2009; Angilletta, 2009) posits that TPCs evolve under severe constraints, due to the impact of thermodynamics on enzyme kinetics. More precisely, it predicts that a rise in the peak temperature (*T*_pk_) through adaptation to a warmer environment will necessarily lead to an increase in the maximum height of the curve (*B*_pk_; Fig. 2A). This adaptive increase in *B*_pk_ with temperature is assumed to be the direct outcome of the acceleration of enzyme-catalysed reactions because enzyme activity also has a unimodal relationship with temperature (Feller, 2010). Hotter-is-better is implicit in the “Universal Temperature Dependence” (UTD) concept of the Metabolic Theory of Ecology (MTE; Brown et al. 2004). MTE also (implicitly) posits that increases in *B*_pk_ should be associated with body size declines, as metabolic trait performance (normalised to a standard temperature) scales negatively with body size across diverse taxonomic groups (Brown et al., 2004). In its strictest form (Fig. 2A), hotter-is-better makes a number of strong and possibly unrealistic assumptions. First, because of thermodynamic constraints, *E* (a measure of thermal sensitivity, also referred to as “activation energy”; Fig. 1) is expected to vary very little across species, with negligible capacity for environmental adaptation (the UTD; Gillooly et al. 2006). Second, if *B*_0_ is calculated at a low enough normalisation temperature (*T*_ref_), then it is expected to be nearly invariant, that is, performance at a low temperature would be almost constant across species (Fig. 2A). This also implies that the body size-scaling of growth rate predicted by MTE must occur at temperatures close to the peak of the curve and not at a low *T*_ref_.

Thus, under the strict hotter-is-better hypothesis, both *B*_0_ (at a low *T*_ref_) and *E* must be nearly constant across species. Relaxing at least one of these leads to a more realistic weak hotter-is-better hypothesis (Figs. 2B; also see SI Fig. S1 for three variants). For example, variation in *B*_0_ can arise from vertical shifts in the whole TPC, driven by changes in body size (Brown et al., 2004). Part of the variation in *B*_0_ may also be driven by the process of metabolic cold adaptation, which results in an elevation of baseline performance in species adapted to low-temperature environments (e.g., see Wohlschlag 1960; Clarke 1993; Seibel et al. 2007; White et al. 2012; Clarke 2017; DeLong et al. 2018). Similarly, recent work has shown that significant variation in *E* exists within and across species, suggesting that this variation is likely adaptive (Dell et al., 2011; Nilsson-Örtman et al., 2013; Pawar et al., 2016; García-Carreras et al., 2018). In any case, under a weak hotter-is-better hypothesis, growth rate is still expected to increase with temperature but the correlation between *T*_pk_ and *B*_pk_ should be weaker.

A third hypothesis, also lying in the middle of the spectrum, is the specialist-generalist tradeoff hypothesis (Huey and Hertz 1984; Angilletta 2009; Fig. 2C). It posits a tradeoff between maximum trait performance (*B*_pk_) and thermal niche width (*W*_op_). That is, a widening of the niche necessarily incurs a metabolic cost (e.g., a cost in enzyme performance), leading to a decrease in peak performance. Note that the weak hotter-is-better and the specialist-generalist tradeoff hypotheses are not mutually exclusive, as their predictions stem from very different mechanisms which could potentially interact.

Finally, at the other end of the spectrum lies a class of hypotheses which posit that the influence of thermodynamic constraints should be minimised through adaptation of species’ biochemical machinery (Hochachka and Somero, 2002; Clarke and Fraser, 2004; Angilletta, 2009; Clarke, 2017). An extreme example is “perfect biochemical adaptation”, which posits that adaptation should allow species to maximise their performance (*B*_pk_) in any thermal environment (Fig. 2D) by overcoming biochemical constraints. An upper limit to the maximum possible *B*_pk_ across species or evolutionary lineages would still exist, but due to a different thermodynamic constraint from that expected under hotter-is-better. This hypothesis further predicts the existence of strong correlations between environmental conditions and TPC parameter values (due to adaptation), with the exception of *B*_pk_ which would be nearly invariant. While some studies have found support for biochemical adaptation (e.g., for photosynthesis rate; Padfield et al. 2016, 2017), it remains unclear whether adaptation can indeed equalize *B*_pk_ across different environments.

The above hypotheses are not an exhaustive list but lie on a spectrum (Fig. 2). To understand the position of different metabolic traits and/or species groups on this spectrum, it is necessary to investigate i) the correlations between multiple thermal parameters and ii) how each thermal parameter evolves across species. Here we tackle this challenge by performing a thorough phylogenetic analysis to investigate the evolution of TPCs of a fundamental measure of fitness—population growth rate (*r*_max_)—using a global database for phytoplankton species. We chose phytoplankton as a study system for ecological and practical reasons. First, phytoplankton form the autotroph base of most aquatic food webs and contribute around half of the global primary production (Field et al., 1998). Second, phytoplankton are one of the few species groups for which sufficiently large TPC datasets for growth rate are available.

Within phytoplankton, we also explore whether the impact of thermodynamic constraints on the shape of the TPC varies between freshwater and marine species. In particular, as freshwater phytoplankton have a relatively more limited potential for dispersal compared to marine phytoplankton which are passively moved by ocean currents across large distances (Doblin and van Sebille, 2016), the timescale of temperature fluctuations that the former experience can be quite different from that of the latter. Such intricacies of the marine environment could potentially be reflected in the TPCs of marine species, which could be under stronger selection for overcoming thermodynamic constraints. Through this detailed analysis, we also shed light on the processes that underlie the Eppley curve (i.e., the relationship between the maximum possible growth rate of all marine phytoplankton and temperature; Eppley 1972), which is widely used in marine ecosystem models (e.g., Palmer and Totterdell 2001; Stock et al. 2014).

## Methods

To understand whether and how thermodynamic constraints influence the evolution of the shape of TPCs of phytoplankton, here we analyse the correlations between TPC parameters across species. For this, we take a phylogenetic comparative approach which allows us to partition the covariance between six TPC parameters of phytoplankton into a phylogenetically heritable component, a fixed effects component, and a residual component. To this end, we estimate the amount of heritable covariance by building a phylogeny of the species in our dataset and combining it with multi-response regression models. To reduce confounding effects of the local environment on TPC parameter covariances, we control for the habitat of species/strains as well as the latitude of their isolation locations through the fixed effects component of our models. For marine species in particular, we also simulate the trajectories of drifting marine phytoplankton to get realistic estimates of the temperatures that they experience through drifting.

### Data

We compiled a global database on growth rate performance of phytoplankton species by combining the previously published datasets of López-Urrutia et al. (2006), Rose and Caron (2007), Bissinger et al. (2008), and Thomas et al. (2012). Growth rates across temperatures were typically measured under light- and nutrient-saturated conditions in these studies. Species names were standardised by querying the Encyclopedia of Life (Parr et al., 2014) via the Global Names Resolver (Global Names Architecture, 2017), followed by manual inspection. This ensured that synonymous species names were represented under a common name. From 795 original species/strain names, this process yielded 380 unique taxa from nine phyla. Where multiple strains of the same species (or isolates from different locations) were available, we did not perform any averaging of growth rate measurements, but analyzed each isolate separately. This allowed us to capture both the inter- and intraspecific variation, where possible. The isolation locations of species/strains in the dataset ranged in latitude from 78°S to 80°N (Fig. S2 in the Supporting Information (SI)).

For cell volume data, those available from original studies were combined with median volume measurements reported by Kremer et al. (2014) and with measurements from Kremer et al. (2017). This process resulted in a dataset with cell volumes for 132 species of phytoplankton, spanning seven orders of magnitude.

### Estimation of TPC parameter values

To quantify all key features of the shape of each growth rate TPC, we used a modified formulation (with *T*_pk_ as an explicit parameter; SI section S3.1) of the four-parameter variant of the Sharpe-Schoolfield model (Schoolfield et al. 1981; Fig. 1):

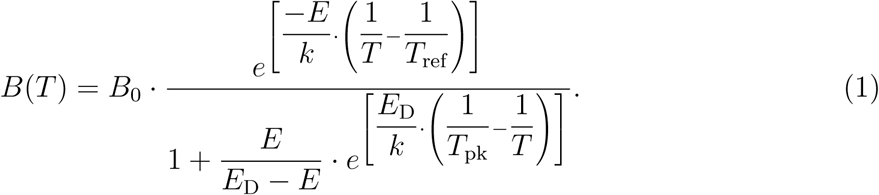

Here, the growth rate, *B* (s^−1^), at a given temperature *T* (K) is expressed as a function of four parameters (*B*_0_, *E, E*_D_ and *T*_pk_; see Fig. 1 for their description and units), and the Boltzmann constant, *k* (8.617·10^−5^ eV · K^−1^). The key assumption of this model is that growth rate is controlled by a single rate-limiting enzyme which is deactivated at high temperatures, and which operates at a decreased rate at low temperatures because of low available kinetic energy. While this assumption is shared by many TPC models, its validity remains under debate (Clarke 2017; DeLong et al. 2017). For example, growth rate may be determined by the effects of temperature on both the activity and the stability (free energy) of one or multiple catalyzing enzymes (DeLong et al., 2017). Other factors besides enzyme thermodynamics might also be important, such as the transport of reaction products in the cell (Ritchie, 2018). Nevertheless, the Sharpe-Schoolfield model remains widely used because it adequately captures the relationship between metabolic traits and temperature (e.g., see Padfield et al. 2016; Salis et al. 2016; Bestion et al. 2018; Francis et al. 2019).

Furthermore, because the Sharpe-Schoolfield model has an exponential term in its numerator (Eq. (1)), *B*(*T*) values estimated with the model will necessarily be positive. Therefore, any negative or zero growth rate measurements had to be removed from the dataset before fitting the model to data. As we show in section S3.2 of the SI, using a model that can accommodate non-positive growth rates instead of the Sharpe-Schoolfield model would not qualitatively change the results of our study. Thereafter, we fitted the Sharpe-Schoolfield model separately to each species/strain in the dataset, using the Levenberg-Marquardt non-linear least squares minimization algorithm (SI section S3.3). After obtaining estimates of the four main model parameters, we used them to calculate the values of two more parameters: *B*_pk_ and *W*_op_ (K) (Fig. 1). We note that we focus on *W*_op_ and not the full niche width of the TPC as i) most species typically experience temperatures well below *T*_pk_ (Martin and Huey, 2008; Thomas et al., 2012; Pawar et al., 2016), and ii) experimentally-determined TPCs typically do not cover a sufficient temperature range to estimate the full niche width (Dell et al., 2011; Pawar et al., 2016).

For a correct comparison of *B*_0_ estimates, *T*_ref_ needs to be set lower than the minimum *T*_pk_ in the dataset. Otherwise, for certain TPCs, *B*_0_ is estimated at the fall of the curve instead of the rise, and the comparison becomes meaningless. As there were a few fits with *T*_pk_ values close to 0°C, we set *T*_ref_ to 0°C (i.e., 273.15 K). However, to ensure that a performance comparison at 0°C does not bias the results of this study—given that some species may not tolerate that low a temperature—, we also fitted the Sharpe-Schoolfield model using a *T*_ref_ of 10°C (i.e., 283.15 K). In that case, we excluded fits with *T*_pk_ < 10°C. All subsequent analyses were performed using both datasets (i.e., those obtained with a *T*_ref_ of 0°C and 10°C), to identify potential areas of disagreement. Finally, as the estimate of *B*_0_ from the Sharpe-Schoolfield model is an approximate measure of the TPC value at *T*_ref_ (*B*(*T*_ref_)) and can sometimes strongly deviate from it, depending—among others—on the choice of *T*_ref_ (Kontopoulos et al., 2018), we calculated *B*(*T*_ref_) manually after obtaining estimates of the four main model parameters (we henceforth refer to *B*(*T*_ref_) as *B*_0_).

Quality filtering of the fits resulted in a TPC dataset of 270 curves using a *T*_ref_ of 0°C and of 259 curves using a *T*_ref_ of 10°C (SI Figs. S4 and S5).

### Reconstruction of the phytoplankton phylogeny

We reconstructed the phylogeny of the species in our TPC dataset using nucleotide sequences of the small subunit rRNA gene (see Table S21 in the SI). One sequence was collected per species where possible, resulting in a dataset of 138 nucleotide sequences. Given that increased taxon sampling has been shown to improve the quality of phylogenetic trees (Nabhan and Sarkar, 2012; Wiens and Tiu, 2012), we also collated a second dataset of 323 sequences by expanding the previous dataset with further sequences of phytoplankton, macroalgae, and land plants. The two sets of nucleotide sequences were aligned with MAFFT (v7.123b; Katoh and Standley 2013), using the L-INS-i algorithm. We then used the entire alignments to build phylogenetic trees without masking any columns, as this has been shown to occasionally result in worse topologies when only a single gene is used (Tan et al., 2015).

Tree topologies were inferred with RAxML (v. 8.2.4; Stamatakis 2014), PhyML (v. 20151210; Guindon et al. 2010), and ExaBayes (v. 1.4.1; Aberer et al. 2014), under the General Time-Reversible model (Tavaré, 1986) with Γ-distributed rate variation among sites (four discrete rate categories; Gu et al. 1995). For RAxML, in particular, we inferred 300 distinct topologies using the slow hill-climbing algorithm (which performs a more thorough exploration of likelihood space than the default algorithm; option “`-f o`”), and selected the tree topology with the highest log-likelihood. For PhyML we used the default options, with the exception of the topology search which was set to include both the Nearest Neighbor Interchange (NNI) and the Subtree Pruning and Regrafting (SPR) procedures. For ExaBayes, we executed four independent runs with four Metropolis-coupled chains per run for 55 million generations. Samples from the posterior distribution were obtained every 500 generations, after discarding the first 25% of samples as burn-in. We confirmed that the four ExaBayes runs had converged through a range of tests (see sections S4.1 and S4.2 in the SI), and obtained a tree topology by computing the extended majority-rule consensus tree. The best tree topology—among those produced by RAxML, PhyML, and ExaBayes—was selected on the basis of proximity to the Open Tree of Life (Hinchliff et al., 2015), and log-likelihood (SI section S4.3).

We then estimated relative ages for all nodes of the best topology, using the uncorrelated Γ-distributed rates model (Drummond et al., 2006), as implemented in DPPDiv (Heath et al., 2012; Flouri and Stamatakis, 2012). To this end, we executed five independent runs for 750,000 generations, sampling from the posterior distribution every 100 generations. As before, we discarded the first 25% of samples as burn-in, and performed diagnostic tests to ensure that the posterior distributions of the four runs had converged and that the parameters were adequately sampled (SI section S4.4). To obtain the final relative time-calibrated tree, we sampled every 300th tree from each run (after the burnin phase) for a total of 9,375 trees, and calculated the median age estimate for each node using the TreeAnnotator program (Rambaut and Drummond, 2017).

### Modelling the local thermal environments of marine phytoplankton

As mentioned previously, although marine phytoplankton are passively moved by ocean currents across large distances, little attention has been given to the potential effects of this on their thermal physiology. In particular, Doblin and van Sebille (2016) showed that the temperature range that marine microbes likely experience is usually much wider if oceanic drifting is properly accounted for. Therefore, to accurately quantify the thermal regimes of marine phytoplankton, we simulated Lagrangian (drifting) trajectories with the Python package OceanParcels (Lange and van Sebille, 2017). More precisely, we used hydrodynamic data from the OFES model (ocean model for the Earth Simulator; Masumoto et al. 2004) to estimate 3,770 backwards-in-time replicate trajectories for each marine location in the dataset over 500 days (using a one-day timestep), at a depth of 2.5, 50, or 100 meters (where possible). These depth values were chosen after considering global estimates of oceanic euphotic depth (Morel et al., 2007), i.e. the depth below which net primary production by marine autotrophs becomes negative (Falkowski and Raven, 2013).

We then calculated the following environmental variables: i) the median temperature experienced, ii) the median latitude visited, iii) the interquartile range of temperatures, and iv) the interquartile range of latitudes. The median captures the central tendency of the temperatures or latitudes that phytoplankton experienced, whereas the interquartile range is a measure of deviation from the central tendency. Measuring all four variables is important, as each of them may have a different effect on the shape of the TPC. The values of the variables were first calculated for each trajectory over the full duration of 500 days, but also over the first 350, 250, 150, and 50 days. They were then averaged across all replicate trajectories per location, depth, and duration, weighted by the length of the trajectory, as some trajectories could be estimated for fewer than 500 days. These variables are hereafter referred to as 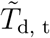 (median temperature), 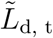 (median latitude), IQR(*T*_d, t_) (interquartile range of temperatures), and IQR(*L*_d, t_) (interquartile range of latitudes), where d and t stand for the depth and duration of the trajectory respectively.

We also obtained temperature data of the isolation locations of marine phytoplankton, in order to compare their explanatory power with that of the Lagrangian trajectory variables. To this end, we used the NOAA Optimum Interpolation Sea Surface Temperature dataset, which comprises daily measurements of sea surface temperature at a global scale and at a resolution of 1/4° (Banzon et al., 2016). Currently, two variants of this dataset are available: i) “AVHRR-Only” which is primarily based on the Advanced Very High Resolution Radiometer, and ii) “AVHRR+AMSR” which also uses data from the Advanced Microwave Scanning Radiometer on the Earth Observing System. The latter variant is considered more accurate, but, for technical reasons, is only available from 2002 until 2011, whereas the former variant is available from 1981 until the present day. In our case, we obtained a daily sea surface temperature dataset between the 1st of September 1981 and the 25th of June 2017, using AVHRR-Only, or AVHRR+AMSR when that was available. From this dataset, we calculated the median temperature of each marine location 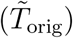, and the interquartile range of temperatures (IQR(*T*_orig_)).

### Inference of TPC parameter co-evolution and associations with environmental variables

#### I. Across the entire dataset

To infer the interspecific correlation structure among the parameters of the TPC and simultaneously detect associations with the local environment of the species in our study, we fitted phylogenetic Markov Chain Monte Carlo generalised linear mixed models using the R package MCMCglmm (v. 2.24; Hadfield 2010). This package can be used to fit phylogenetic regression models, enabling the partitioning of phenotypic trait variance into a phylogenetically heritable component, a fixed effects component of explanatory variables, and a residual variance component (i.e., variance that should be mostly due to environmental effects that are not already controlled for). For the purposes of this study, we constructed multi-response regression models (i.e., models with multiple response variables instead of one), in which the response comprised all six TPC parameters. In other words, instead of trying to predict a single response variable, the models would predict all six TPC parameters, while simultaneously inferring their variance/covariance matrix. Each element of this matrix was estimated as a free parameter from the data, so that any correlations between pairs of TPC parameters could be detected.

To ensure that the distribution of each response variable was as close to normality as possible, we applied a different transformation to each TPC parameter: 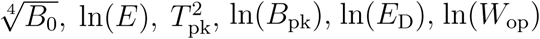. It was necessary to perform those transformations as each response variable in an MCMCglmm needs to conform to one of the implemented distributions in the package (e.g., Gaussian, Poisson, multinomial), with the Gaussian distribution being the most appropriate here. Besides this, most macroevolutionary models assume that the evolutionary change in trait values follows a Gaussian distribution. Thus, statistical transformations of trait values are often used to satisfy this assumption. In any case, applying these transformations does not affect our results qualitatively even though thermal parameter correlations are estimated in transformed (not linear) scale. To incorporate the uncertainty for each transformed thermal parameter estimate, we used the delta method (e.g., see Oehlert 1992) implemented in the R package msm (v. 1.6.4; Jackson 2011) to obtain appropriate estimates of the variance of the standard error for 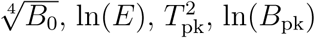, and ln(*E*_D_). As we manually calculated ln(*W*_op_) a posteriori without an analytical solution, we performed bootstrapping to obtain error estimates for it.

For the majority of the TPCs in our dataset, there was at least one parameter whose value could not be estimated with certainty due to lack of adequate experimental measurements (SI section S3.3). MCMCglmm can accommodate such missing values in the response by treating them as “Missing At Random” (MAR; see Hadfield 2010, de Villemereuil and Nakagawa 2014, and Tierney and Cook 2018). The MAR assumption is valid as long as i) missing values in a response variable can be estimated (with some uncertainty) from other components of the model (i.e., other covarying response variables or the phylogeny), and ii) data missingness is not driven by a variable that is not included in the model. When these two conditions are true, the inferred estimates of missing values are unbiased (see Nakagawa and Freckleton 2008; Garamszegi and Møller 2011). Applying this method allowed us to include TPC parameter estimates from curves that were only partly well sampled (e.g., only the rise of the curve), increasing the statistical power of the analysis and reducing the possibility of estimation biases (e.g., in the covariances among TPC parameters).

The fixed effects component of each candidate model contained at the very minimum a distinct intercept for each response variable. Starting with this, we fitted models with i) no other predictors (the intercepts-only model), ii) the latitude of the isolation location of each species, iii) the habitat of each species (marine vs freshwater), or iv) both latitude and habitat. For models that included latitude as a predictor, we specified either the absolute latitude of the location or a second order polynomial (because mean environmental temperature and its fluctuations are approximately unimodal functions of latitude from the equator to mid-latitudes). In any case, we estimated the association of each fixed effect (latitude and/or habitat) with each response variable separately (by inferring distinct coefficients for, e.g., ln(*E*):|latitude|, ln(*B*_pk_):|latitude|). It is worth noting that we did not include the temperature of the environment as a fixed effect in these particular models, as there was no reliable temperature dataset with high enough resolution for both marine and freshwater locations. To avoid any potential biases introduced by a combination of two temperature datasets (one for the marine locations and one for the freshwater ones), we instead used latitude as a proxy for temperature variation.

Species identity was specified as a random effect on the intercepts. To integrate phylogenetic information into the model, we first pruned the phylogeny to the subset of species for which data were available (SI Fig. S15). We next calculated the inverse of the phylogenetic covariance matrix from the phylogenetic tree, including ancestral nodes as this allows for more computationally efficient calculations (Hadfield and Nakagawa, 2010; de Villemereuil and Nakagawa, 2014).

The default prior was used for the fixed effects, whereas for the random effect and the residual variance components, we used a relatively uninformative inverse-Γ prior with shape and scale equal to 0.001 (the lower this number the less informative is the prior). For each model, two chains were run for 100 million generations, sampling from the posterior distribution every 1000 generations after discarding the first 10 million generations as burnin. Convergence between each pair of chains was verified by calculating the potential scale reduction factor (Gelman and Rubin, 1992; Brooks and Gelman, 1998) for all estimated parameters (i.e., fixed effects, elements of the phylogenetically heritable and residual matrices), and ensuring that it was always lower than 1.1. We also confirmed that the effective sample size of all model parameters—after merging samples from the two chains—was greater than 200, so that the mean could be adequately estimated.

Model selection was done on the basis of the Deviance Information Criterion (DIC; Spiegelhalter et al. 2002), averaged across each pair of chains. We excluded models if a fixed effect had a 95% Highest Posterior Density (HPD) interval that included zero for every single response variable (e.g., if all of 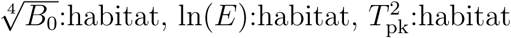 etc. had 95% HPD intervals that included zero). In frequentist statistics terms, this is roughly equivalent to excluding models whose predictors were not significant for any response variable.

Phenotypic correlations between pairs of TPC parameters (*r*_phe_) were broken down into their phylogenetically heritable (*r*_her_) and residual components (*r*_res_) by dividing the covariance estimate between two parameters by the geometric mean of their variances. These were inferred from the best-fitting model in terms of DIC.

Finally, we measured the phylogenetic heritability (i.e., the ratio of heritable variance to the sum of heritable and residual variance) of each TPC parameter. As the phylogeny is integrated with the MCMCglmm, the resulting estimates are equivalent to Pagel’s *λ* (Pagel, 1999; Hadfield and Nakagawa, 2010), and reflect the strength of the phylogenetic signal, i.e., the extent to which closely related species are more similar to each other than to any species chosen at random (Pagel, 1999; Kamilar and Cooper, 2013; Symonds and Blomberg, 2014). Strong phylogenetic signal would indicate that variation in the TPC parameter can be explained by its gradual evolution across the phylogeny. On the other hand, a lack of phylogenetic signal would reflect either trait stasis (with any variation among species being noise-like) or very rapid evolution (that is independent of the phylogeny). Intermediate values of phylogenetic signal would imply either that the TPC parameter is under constrained evolution (e.g., due to stabilizing selection), or that its evolutionary rate changes through time (e.g., an evolutionary rate acceleration could lead to the convergence of the niches of distantly related species). We obtained phylogenetic heritability estimates from the intercepts-only model as the addition of fixed effects would reduce the residual variance and bias the heritability estimates towards higher values.

#### II. For the marine subset of the data

To test whether the correlation structure of thermal parameters across the entire phytoplankton dataset differs from that of marine species only, we also performed the above analysis for only the marine species in the dataset. The main difference in the specification of the MCMCglmms for marine species was that we used fixed effects that captured both the latitude and the temperature characteristics of the local environment of phytoplankton (see the “Modelling the local thermal environments of marine phytoplankton” section above): i) no fixed effects (intercepts-only model), ii) *L*_orig_, iii) 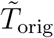, iv) IQR(*T*_orig_), v) 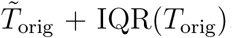, vi) 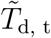, vii) IQR(*T*_d, t_), viii) 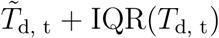, ix) 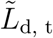, x) IQR(*L*_d, t_), xi) 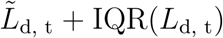. All latitude variables other than IQR(*L*_d, t_) were specified—in different models—both as a second order polynomial and with absolute values. A second order polynomial was also tested for IQR(*T*_d, t_) variables to investigate the existence of a quadratic relationship of IQR(*T*_d, t_) with thermal parameters.

As there was a very large number of MCMCglmms to execute (158 pairs of chains), we first ran each of them for 60 million generations. We then checked whether the two chains per model had converged as previously described, and reran the subset that had not converged for 120 million generations. At that point, all pairs of chains converged on statistically indistinguishable posterior distributions. As above, samples from the first 10% generations of each model were discarded as burn-in.

### Size-scaling of *B*_0_ and *B*_pk_

As explained in the introduction, MTE predicts that temperature-normalised *r*_max_ should be constrained by body mass across taxonomic groups (Brown et al., 2004). However, at finer taxonomic resolutions (e.g., within species), this relationship may take the opposite direction, i.e., selection for high *r*_max_ may lead to declines in body size as has been observed widely (the “temperature-size rule”; Atkinson 1996; Winder et al. 2009; Yvon-Durocher et al. 2011; Peter and Sommer 2013; Sommer et al. 2017). For example, warming may confer a competitive advantage to smaller phytoplankton due to their higher *r*_max_ (Reuman et al., 2014). Therefore, as a final step for understanding how TPCs evolve, we tested whether and how growth rate scales with body size. Under the strict hotter-is-better hypothesis, such scaling would be expected only for growth rates near each species’ *T*_pk_, whereas if the weak hotter-is-better hypothesis holds, size scaling could also—but not necessarily—occur at low temperatures. Understanding if the latter holds requires first the calculation of growth rate values for all TPCs at a common normalisation temperature (*T*_ref_), followed by their examination for any size scaling patterns. Therefore, to test both hypotheses of body size-scaling, we fitted MCMCglmms with cell volume as a fixed effect and a single response of either i) *B*_0_ (at a *T*_ref_ of 0°C), ii) *B*_0_ (at a *T*_ref_ of 10°C), or iii) *B*_pk_. Species identity was treated as a random effect on the intercept, the slope, or both. Each model was fitted with and without the phylogenetic variance/covariance matrix to compare the predictions obtained by ignoring the phylogeny or accounting for it. Two chains were run per model for a length of 3 million generations, and convergence was established as in the previous section after removing samples from the first 300,000 generations. DIC was used to identify the most appropriate model for each response variable. To evaluate the quality of the best-fitting model, we first calculated the amounts of variance explained by fixed 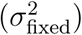 and random effects 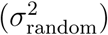, and the residual variance 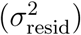. From these, we calculated the marginal 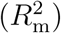 and conditional 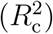 coefficients of determination, as described by Nakagawa and Schielzeth (2013):

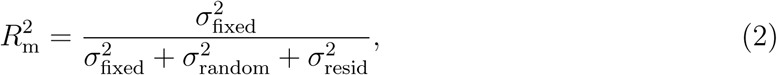

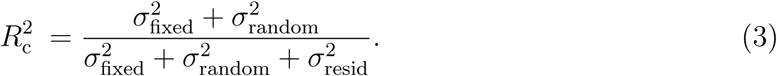

## Results

### Interspecific correlations and phylogenetic signal

The best-fitting phylogenetic regression model on the basis of DIC had only latitude as a fixed effect (SI Fig. S16). Models with habitat as a predictor were excluded from the DIC comparison, as the 95% Highest Posterior Density interval of every single habitat coefficient included zero. This likely reflects that any effects of habitat type on TPC parameters are already captured by the phylogenetic correction, especially given that phytoplankton habitat is phylogenetically structured (SI Fig. S15). In contrast, the 95% HPD intervals of the coefficients of latitude for *T*_pk_ (for both *T*_ref_ values) and *E* (for a *T*_ref_ of 0°C only) did not include zero (SI Fig. S17). A minor difference between the analyses with a *T*_ref_ of 0°C and 10°C was that in the former case, the model with a second order polynomial in latitude was selected, whereas in the latter case, absolute latitude performed better. The shapes of the two fitted curves (SI Fig. S17) suggest that the effect of latitude on the TPC is particularly strong for colder-adapted species, leading to a deviation from a strictly linear association.

From the analysis of the resulting interspecific variance/covariance matrices (SI Tables S4, S5, S8, and S9), we identified only two correlations among TPC parameters: i) between *B*_pk_ and *T*_pk_ (Fig. 3A), and ii) between *E* and *W*_op_ (SI Fig. S18). The former correlation appears to be driven entirely by the phylogenetically heritable (*r*_her_) component of the coldest-adapted species in the dataset (i.e., the three data points with *T*_pk_ < 10°C in Fig. 3A), and becomes nonexistent when these are excluded. Such a weak correlation is consistent with the weak hotter-is-better hypothesis (Fig. 2). Also, as *E* and *W*_op_ are both measures of thermal sensitivity in the range of temperatures where organisms typically operate, a negative correlation between them was expected under all TPC evolution hypotheses. In contrast, a negative correlation between *B*_pk_ and *W*_op_, which would be expected by the specialist - generalist tradeoff hypothesis, was not supported by the data (Fig. 3B). Finally, we detected varying amounts of phylogenetic signal in all TPC parameters, with *T*_pk_ showing the strongest (perfect phylogenetic) signal (Fig. 4). This was in contrast to the assumptions of the strict hotter-is-better and the perfect biochemical adaptation hypotheses, which posit that *E* and *B*_pk_ respectively should vary very little across species and not in a phylogenetically heritable manner (Fig. 2).

**Figure 3.**
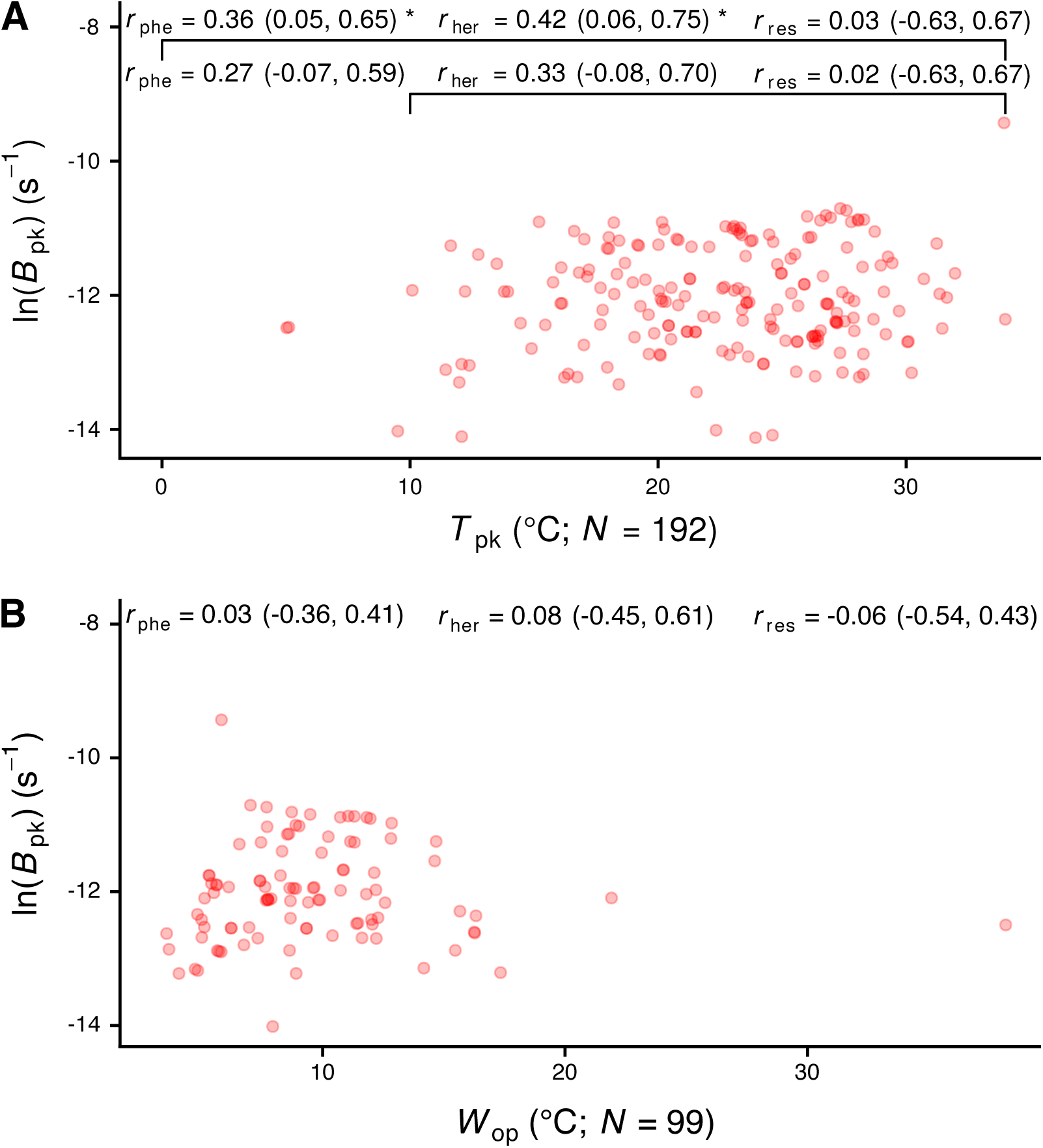
The relationship of *B*_pk_ with *T*_pk_ (A) and *W*_op_ (B). *r*_phe_, *r*_her_, and *r*_res_ stand for phenotypic correlation, phylogenetically heritable correlation and residual correlation respectively. The three correlation coefficients were simultaneously inferred after correcting for phylogeny and for environmental effects (latitude). For panel A, in particular, correlations were estimated across the entire dataset, and after excluding data points with *T*_pk_ < 10°C. The reported estimates are for the correlation of ln(*B*_pk_) with 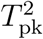 and ln(*W*_op_) respectively, but the horizontal axes are shown in linear scale for simplicity. Values in parentheses correspond to the 95% HPD interval of each correlation coefficient.

**Figure 4.**
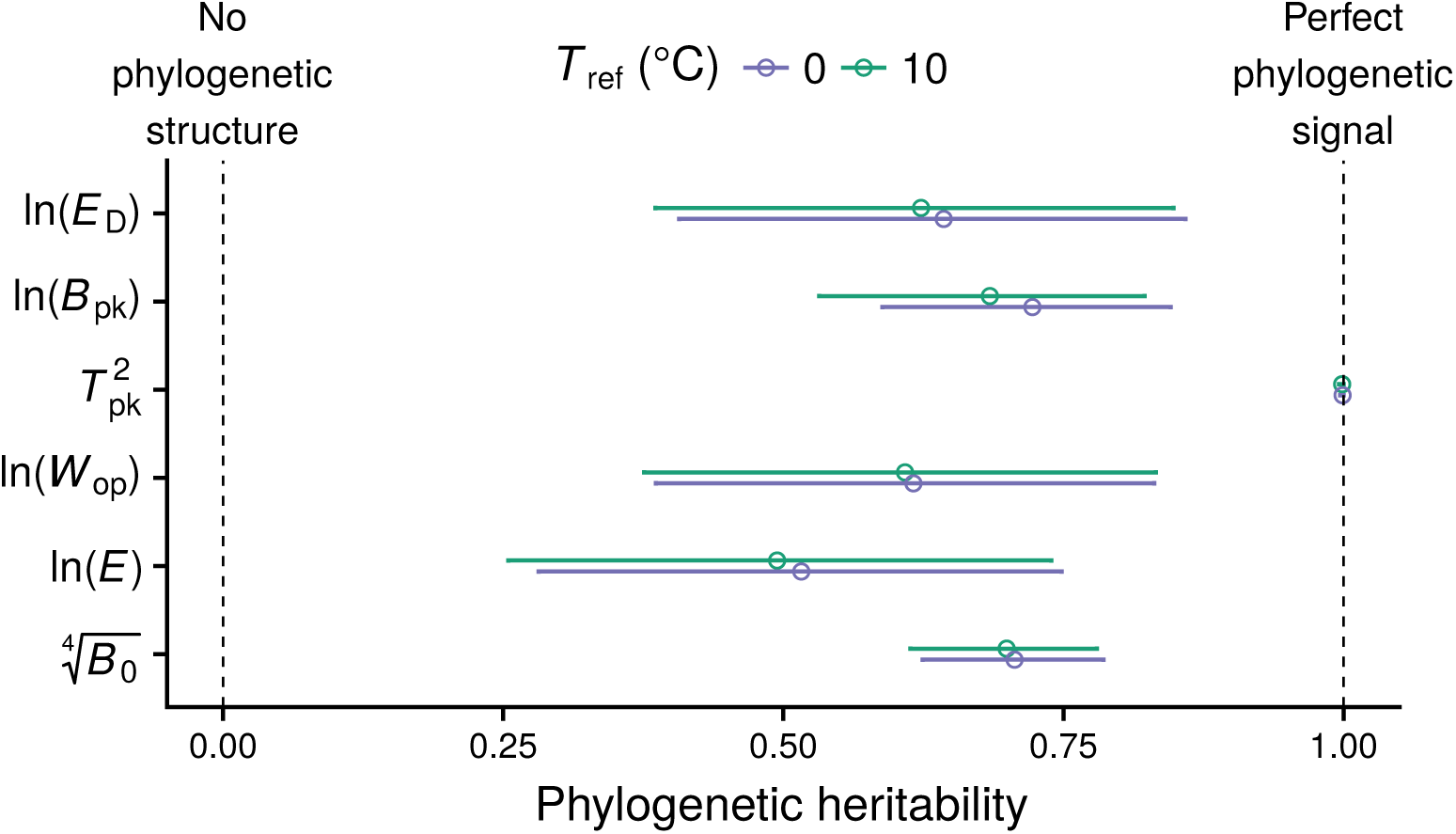
Phylogenetic heritability estimates across the TPC. Circles indicate the mean of the posterior distribution, whereas horizontal bars show the 95% HPD interval. Note that each TPC parameter was transformed towards approximate normality in order to satisfy the requirement of the MCMCglmm method.

Running MCMCglmms for the marine species only yielded mostly similar conclusions (SI section S5.2). The only correlation that could be detected was between *E* and *W*_op_ (SI Fig. S22). The best-fitting model had a fixed effect of 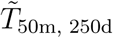 (for *T*_ref_ = 0°C) or IQR(*T*_50m, 50d_) (for *T*_ref_ = 10°C). More precisely, the analysis of all marine species revealed a negative relationship between ln(*B*_pk_) and the median temperature of trajectories at a depth of 50 meters and for a duration of 250 days (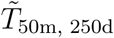; SI Fig. S20). If, instead, only marine species with *T*_pk_ > 10°C are included, ln(*E*) is the parameter that associates with the environment, increasing with the interquartile range of temperatures of trajectories at a depth of 50 meters and for a duration of 50 days (IQR(*T*_50m, 50d_); SI Fig. S21). For both *T*_ref_ values, the best-fitting Lagrangian models had consistently lower DIC values (at least 30 DIC units difference; see SI Tables S10 and S15) than their non-Lagrangian equivalents.

### Size-scaling of growth rate

Cell volume-growth rate scaling as predicted by the MTE and expected by the two (strict and weak) hotter-is-better hypotheses, was detected only in the maximum height of the curve (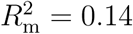 and 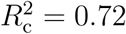; Fig. 5C-D) and not at the performance at a temperature of 0°C or 10°C (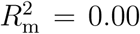 and 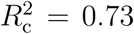; Fig. 5A-B). In particular, across the entire dataset, *B*_pk_ was found to scale with cell volume raised to an exponent of −0.09 (95% HPD interval = (−0.15, −0.05); Fig. 5C). The best-fitting models always had a random effect of species identity on the intercept and not the slope (SI Table S20).

**Figure 5.**
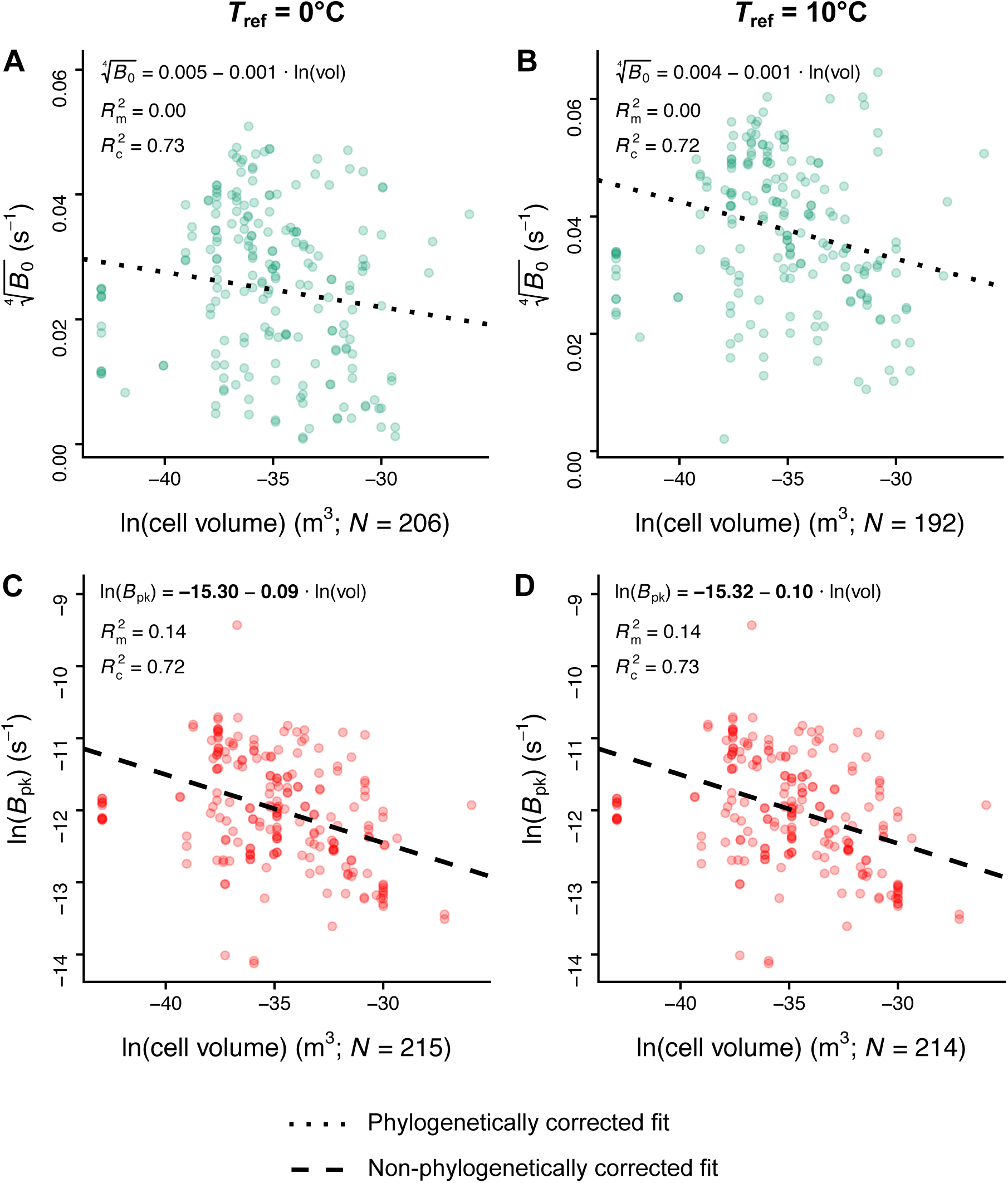
The relationships of cell volume with *B*_0_ (panels A and B) and *B*_pk_ (panels C and D) with *T*_ref_ set to 0°C or 10°C, according to the best-fitting model in each case (see SI Table S20). Coefficients shown in bold had 95% HPD intervals that did not include zero. The sample sizes of *B*_0_ and *B*_pk_ estimates shown here are higher than those reported in SI Figs. S4 and S5, as we included estimates from species with unknown isolation locations. Note that we used different statistical transformations for *B*_0_ and *B*_pk_ so that their estimates would be nearly normally distributed.

## Discussion

In this study we investigated the influence of thermodynamic constraints on the shape of the thermal performance curve of phytoplankton (Fig. 2). To this end, we performed a thorough analysis of correlations among six TPC parameters. Controlling for the phylogeny of species and their local environment allowed us to better tease apart the relationships among thermal parameters and quantify the influence of phylogeny on each TPC parameter.

We detected a positive correlation between *B*_pk_ and *T*_pk_ (Fig. 3A), which was however very weak and only held if TPCs with very low *T*_pk_ values were included. This pattern is inconsistent with the strict hotter-is-better hypothesis (Fig. 2). Therefore, we can conclude that phytoplankton TPCs do not support the strong thermodynamic constraints extreme of the spectrum of hypotheses. The only other correlation that we detected was between *E* and *W*_op_ (SI Fig. S18), which is expected because niche width within the operational temperature range varies inversely with thermal sensitivity (*E*). When focusing only on marine phytoplankton, we detected neither a correlation between *B*_pk_ and *T*_pk_, nor any correlation uniquely present in marine species. However, this may reflect the lower statistical power of the analysis of marine species due to the smaller sample size. In any case, as a correlation between *B*_pk_ and *W*_op_ was not detected in either the analysis of the entire dataset (Fig. 3B) or in the analysis of correlations from marine species, the generalist-specialist tradeoff hypothesis can also be rejected.

To further narrow down the location of phytoplankton TPCs on the spectrum of hypotheses (Fig. 2), we examined the estimates of phylogenetic signal of all six TPC parameters, which were simultaneously inferred from our multi-response regression models. We used the estimates to test both the strict hotter-is-better hypothesis and the perfect biochemical adaptation hypothesis, which predict a complete lack of phylogenetic signal in *E* and *B*_pk_ respectively. This analysis also yielded a basic understanding of how the remaining TPC parameters (e.g., *T*_pk_) evolve. Overall, the mean phylogenetic signal estimate of *E* was the lowest of all TPC parameters, but its 95% HPD interval was well above zero (Fig. 4). This result further supports the rejection of the strict hotter-is-better hypothesis. Moreover, it indicates that *E* is not nearly constant across species—contrary to what the MTE initially assumed (see SI Fig. S1A and Gillooly et al. 2001; Clarke and Fraser 2004; Clarke 2004; Gillooly et al. 2006; Clarke 2006)—, and provides some insight into the inter- and intraspecific variation in *E* reported by previous studies (e.g., Dell et al. 2011; Nilsson-Örtman et al. 2013; Pawar et al. 2016).

At the right end of the hypotheses spectrum, we were also able to reject the perfect biochemical adaptation hypothesis because *B*_pk_ also exhibited phylogenetic signal. It is worth noting that the phylogenetic signal in *B*_pk_ does not merely reflect that the local environment is phylogenetically heritable (with closely related species occurring in geographically close environments), as the correlation between phylogenetic distance and geographical distance was almost zero (SI section S5.1). In any case, variation in *B*_pk_ was not found to be latitudinally structured across the entire dataset (contrary to *E* and *T*_pk_; SI Fig. S17), albeit marine species that experienced low temperatures had slightly higher *B*_pk_ values (SI Fig. S20A). An elevation of *B*_pk_ (or *B*_0_) in organisms living at cold environments could arise from the process of metabolic cold adaptation, which has sometimes been detected in other species groups (see Wohlschlag 1960; Clarke 1993; Seibel et al. 2007; White et al. 2012; Clarke 2017; DeLong et al. 2018).

Lastly, we examined the effect of body size on growth rate. We found a weak negative scaling of *B*_pk_ (maximum height of the TPC) with cell volume, whereas trait values normalised to 0°C or 10°C did not exhibit any size scaling (Fig. 5). This suggests an energetic tradeoff between cell volume and *B*_pk_ in phytoplankton, similar to the prediction of the supply-demand body size optimization model of DeLong (2012). That is, the maintenance of a large cell volume should incur a high energetic cost, reducing the amount of energy that can be directed to cell growth, and vice versa. The weak negative size scaling of *B*_pk_ is consistent with our only remaining hypothesis: the weak hotter-is-better hypothesis. Indeed, given the weak correlations of i) *B*_pk_ with *T*_pk_, and ii) *B*_pk_ with cell volume, an increase in *T*_pk_ would lead to a weak increase in *B*_pk_ and, indirectly, to a weak decrease in cell volume. Therefore, a decrease in cell size with warming—which has often been observed (Winder et al., 2009; Yvon-Durocher et al., 2011; Peter and Sommer, 2013; Sommer et al., 2017)—could be constrained by an indirect correlation between *T*_pk_ and cell volume.

Our results about the weak relationship between *B*_pk_ and *T*_pk_, and the scaling of the former with cell volume are consistent with the conclusions of Kremer et al. (2017). They found evidence for the effects of temperature, taxonomic group, and cell size on the maximum growth rate of phytoplankton, effectively suggesting adaptation of *B*_pk_ across lineages. This further means that the classical Eppley curve (Eppley, 1972; Bissinger et al., 2008) does not necessarily indicate as strong a global (thermodynamic) constraint on maximum performance across species as has been previously thought. In this context, we also note that ideally cell size should be directly accounted for in analyses of TPC evolution. This was partially done in our study (i.e., by examining the relationship of cell volume with *B*_0_ and *B*_pk_), as we could not obtain cell volume measurements for all species in our dataset.

Given all these results, we conclude that the TPCs of phytoplankton evolve in the general absence of hard thermodynamic constraints, similarly to the expectations of a very weak hotter-is-better hypothesis (Fig. 3A). A possible mechanistic interpretation of the observed patterns is that, at very low temperatures, the limiting factor is low available kinetic energy, which constrains the rate of biochemical reactions. At higher temperatures, on the other hand, maximum trait performance appears temperature-independent, suggesting the presence of biophysical or other constraints. For example, given that *B*_pk_ scales negatively with cell volume, a lower limit in cell volume (e.g., due to the need for maintaining non-scalable cellular components such as membranes; Raven 1998) will also set an upper limit to the maximum possible growth rate.

To the best of our knowledge, a thorough analysis of the correlation structure among parameters that control the entire range of the TPC has never been conducted. At most, previous studies have investigated the existence of correlations between two or three selected TPC parameters (e.g., between *T*_pk_ and *B*_pk_; see Frazier et al. 2006 and Sørensen et al. 2018). This can be problematic for two reasons. First, by only focusing on parameters that control the peak of the TPC, such studies ignore potential correlations with parameters in other areas of the curve (e.g., *E*). Second, even if a statistical correlation can be observed between two thermal parameters, it could be driven by the covariance of the two parameters with other, overlooked TPC parameters. Indeed, many studies on TPCs do not explicitly account for phylogenetic relationships among species at all (but see Sal et al. 2015 for a phylogenetically-controlled study on the size-scaling of phytoplankton growth rate). Our results highlight the fact that ignoring potential phylogenetic effects can make it harder to differentiate between alternative hypotheses on the evolution of TPCs, and may leave studies vulnerable to biases introduced by phylogenetic nonindependence (e.g., an observed relationship between two TPC parameters could arise solely from uneven phylogenetic sampling).

Perhaps the most striking result of this study is that we detected a very limited number of correlations or tradeoffs across the entire TPC. One potential explanation for this could be that different phytoplankton lineages have evolved distinct strategies to maximise their fitness. Such strategies may involve thermal parameter correlations that are lineage-specific and hence hard to detect. A similar analysis performed separately for each phytoplankton phylum could potentially address this question. However, obtaining accurate estimates of lineage-specific variance/covariance matrices of TPC parameters would require bigger thermal performance datasets than those that—to our knowledge—are currently available. It would also be interesting to investigate whether the phylogenetic signal of TPC parameters and the correlations among them vary across traits (e.g., photosynthesis rate, respiration rate) or phylogenetic groups (e.g., bacteria, plants). Such analyses could provide useful insights into the nature of possible constraints and their degree of influence on the shape of the thermal performance curve across different branches of the tree of life.

Another direction that could be further pursued involves investigating the effects of the marine environment on phytoplankton TPCs, and, in particular, how TPCs adapt to temperature fluctuations due to oceanic drifting (see e.g., Schaum et al. 2018). It is worth emphasising that, in our study, models that accounted for oceanic drifting of marine phyto-plankton (models with Lagrangian variables) systematically performed better (in terms of DIC) than models that only incorporated the latitude or the sea surface temperature of the isolation locations of the strains. While we detected some associations between environmental variables and TPC parameters, the low sample size and the coarse modelling of drifting prevent us from drawing very strong conclusions. More precisely, some of the limitations of our approach were that simulations were done at only three depths, and did not account for the vertical movement of phytoplankton or the concentration of nutrients. A more in-depth analysis on these matters could be the focus of future studies.

Finally, there is mounting evidence that the shape of TPCs is also affected by a range of other factors such as nutrient availability (Thomas et al., 2017; Bestion et al., 2018), oxygen supply (Gangloff and Telemeco, 2018), and predation risk (Dell et al., 2014; Luhring et al., 2018). Thus, to improve our understanding of how species adapt to different thermal environments, future studies could investigate the adaptive potential of organismal responses not only to temperature, but to the interaction of multiple factors. Such an approach could uncover important adaptive constraints which may not be detectable by studying the responses of biological traits to each factor in isolation.

## Author contributions

DGK and SP conceived and designed the study. DGK performed all analyses other than the simulations of marine phytoplankton trajectories, which were conducted by EvS and ML. DGK, GYD, TGB, and SP interpreted the results. DGK wrote the initial manuscript, with comments and additions provided by all other coauthors.

## Acknowledgements

We thank Bernardo García-Carreras, I. Colin Prentice, Guy Woodward, and Andrew G. Hirst for useful discussions and comments. We are also grateful to the CIPRES Science Gateway (Miller et al., 2010) and Imperial College London’s High Performance Computing service (doi:10.14469/hpc/2232) for access to computational resources. DGK was supported by a Natural Environment Research Council (NERC) Doctoral Training Partnership (DTP) scholarship (NE/L002515/1). SP and GYD were supported by NERC grants NE/M004740/1 and NE/M003205/1 respectively.

