## Supporting Information for "Phytoplankton thermal responses adapt in the absence of hard thermodynamic constraints"

**S6 Scaling of $B_0$ and $B_{pk}$ with cell volume** 33

**S7 List of 16S/18S rRNA sequences used for phylogeny reconstruction** 34

**1.** Science and Solutions for a Changing Planet DTP. **2.** Department of Life Sciences, Imperial College London, Silwood Park, Ascot, Berkshire SL5 7PY, UK. **3.** Grantham Institute, Imperial College London, London SW7 2AZ, UK. **4.** Institute for Marine and Atmospheric Research Utrecht, Utrecht University, Utrecht 3584 CC, the Netherlands. **5.** Department of Earth Science and Engineering, Imperial College London, London SW7 2AZ, UK. **6.** Environment and Sustainability Institute, University of Exeter, Penryn, Cornwall TR10 9EZ, UK. * Corresponding author;.

### S1 Variants of the weak hotter-is-better hypothesis

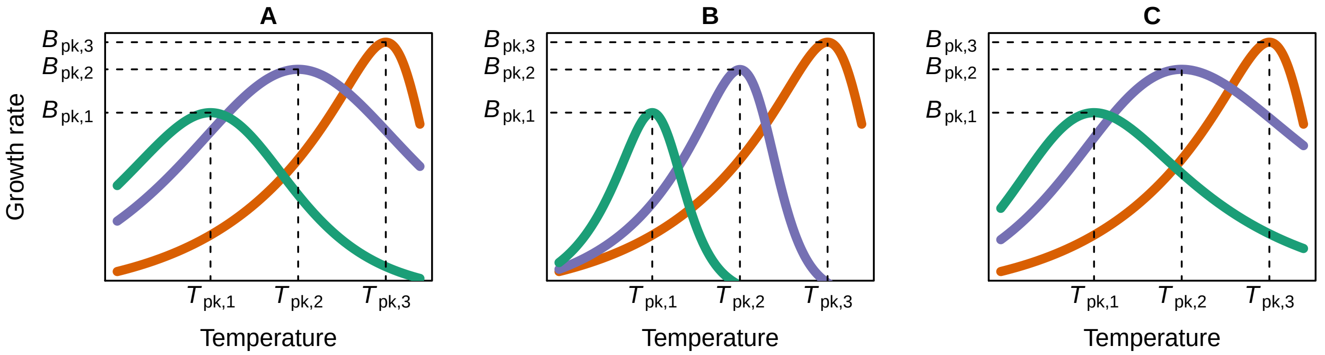

Figure S1: Three alternative ways in which a weak hotter-is-better pattern may emerge. A: The slope of the rising part of the TPC ($E$) remains fixed, whereas the baseline height ($B_0$) varies across species. B: $B_0$ is fixed, whereas $E$ varies. C: Both $B_0$ and $E$ exhibit variation. The variant that is closest to the expectations of the Metabolic Theory of Ecology is that of panel A. To facilitate the comparison of the three variants of weak hotter-is-better, we kept the same $B_{pk}$ and $T_{pk}$ values across the three panels.

### S2 Isolation locations of species/strains in this study

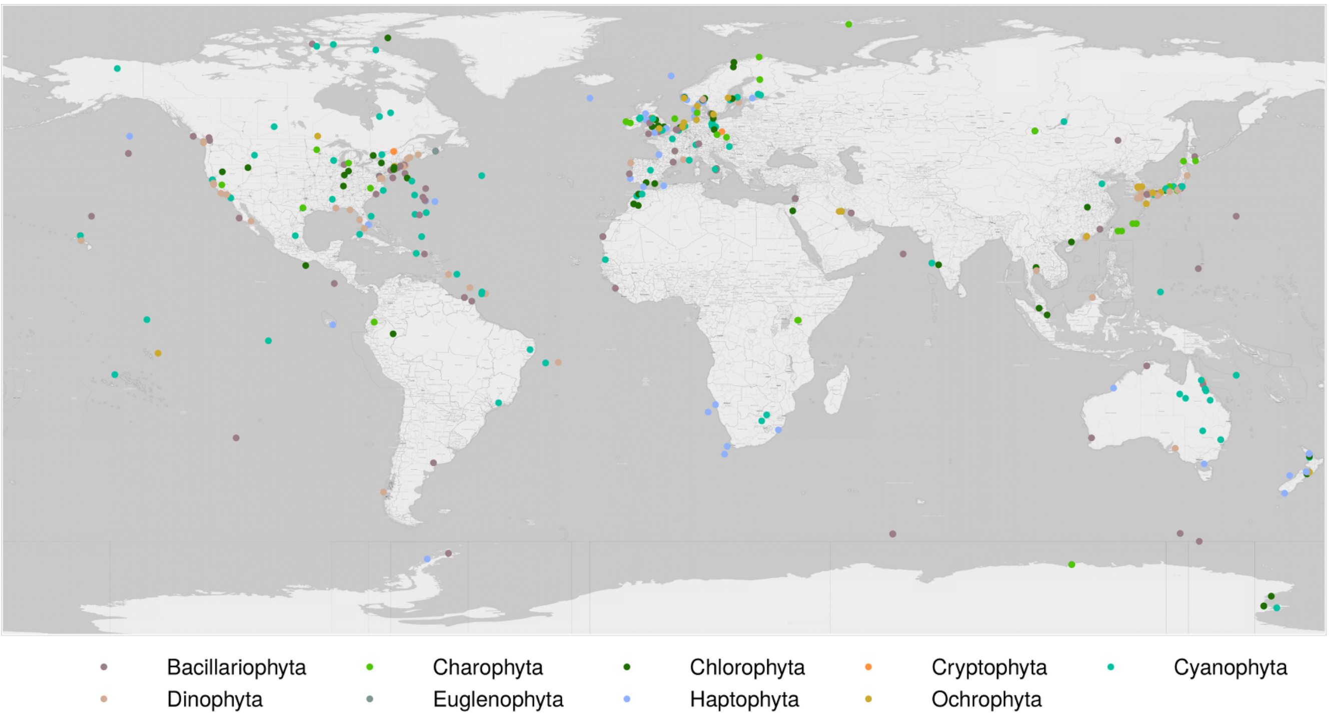

Figure S2: Isolation locations of phytoplankton species, coloured by phylum.

### S3 Estimation of TPC parameter values

#### S3.1 Derivation of the Sharpe-Schoolfield equation with $T_{\mathrm{pk}}$ as a parameter

The original equation of the four-parameter Sharpe-Schoolfield model (Schoolfield et al., 1981) includes the parameter $T_{\mathrm{h}}$, which stands for the temperature at which half of the population of the key rate-limiting enzyme is made inactive:

$$B(T) = B_0 \cdot \frac{e^{\left[\frac{-E}{k}\cdot\left(\frac{1}{T}-\frac{1}{T_{\mathrm{ref}}}\right)\right]}}{1+e^{\left[\frac{E_{\mathrm{D}}}{k}\cdot\left(\frac{1}{T_{\mathrm{h}}}-\frac{1}{T}\right)\right]}}. \tag{S1}$$

As $T_{\mathrm{h}}$ can be lower or higher than the temperature at which the thermal performance curve peaks ($T_{\mathrm{pk}}$), there is not a straightforward mapping between the value of $T_{\mathrm{h}}$ and the effect of temperature on the shape of the thermal performance curve. $T_{\mathrm{pk}}$ is a more useful parameter from a physiological standpoint and, as we had previously shown in Kontopoulos et al. (2018), it can be analytically estimated as

$$T_{\mathrm{pk}} = \frac{-E_{\mathrm{D}} \cdot T_{\mathrm{h}}}{k \cdot T_{\mathrm{h}} \cdot \ln \frac{E}{E_{\mathrm{D}} - E} - E_{\mathrm{D}}}. \tag{S2}$$

Then, to reformulate Eq. (S1) with $T_{\mathrm{pk}}$, we first solve Eq. (S2) for $T_{\mathrm{h}}$:

$$T_{\mathrm{pk}} = \frac{-E_{\mathrm{D}} \cdot T_{\mathrm{h}}}{k \cdot T_{\mathrm{h}} \cdot \ln \frac{E}{E_{\mathrm{D}} - E} - E_{\mathrm{D}}} \implies T_{\mathrm{pk}} \cdot \left(k \cdot T_{\mathrm{h}} \cdot \ln \frac{E}{E_{\mathrm{D}} - E} - E_{\mathrm{D}}\right) = -E_{\mathrm{D}} \cdot T_{\mathrm{h}} \implies$$

$$\implies T_{\mathrm{h}} \cdot \left(k \cdot T_{\mathrm{pk}} \cdot \ln \frac{E}{E_{\mathrm{D}} - E} + E_{\mathrm{D}}\right) = E_{\mathrm{D}} \cdot T_{\mathrm{pk}} \implies T_{\mathrm{h}} = \frac{E_{\mathrm{D}} \cdot T_{\mathrm{pk}}}{k \cdot T_{\mathrm{pk}} \cdot \ln \frac{E}{E_{\mathrm{D}} - E} + E_{\mathrm{D}}}.$$

Finally, we substitute $T_{\mathrm{h}}$ with the above quantity in Eq. (S1):

$$B(T) = B_0 \cdot \frac{e^{\left[\frac{-E}{k}\cdot\left(\frac{1}{T}-\frac{1}{T_{\mathrm{ref}}}\right)\right]}}{1+e^{\left[\frac{E_{\mathrm{D}}}{k}\cdot\left(\frac{1}{T_{\mathrm{h}}}-\frac{1}{T}\right)\right]}} = B_0 \cdot \frac{e^{\left[\frac{-E}{k}\cdot\left(\frac{1}{T}-\frac{1}{T_{\mathrm{ref}}}\right)\right]}}{1+e^{\left[\frac{E_{\mathrm{D}}}{k}\cdot\left(\frac{k \cdot T_{\mathrm{pk}} \cdot \ln \frac{E}{E_{\mathrm{D}} - E} + E_{\mathrm{D}}}{E_{\mathrm{D}} \cdot T_{\mathrm{pk}}}-\frac{1}{T}\right)\right]}} =$$

$$= B_0 \cdot \frac{e^{\left[\frac{-E}{k}\cdot\left(\frac{1}{T}-\frac{1}{T_{\mathrm{ref}}}\right)\right]}}{1+e^{\left[\ln \frac{E}{E_{\mathrm{D}} - E}\right]} \cdot e^{\left[\frac{E_{\mathrm{D}}}{k}\cdot\left(\frac{1}{T_{\mathrm{pk}}}-\frac{1}{T}\right)\right]}} = B_0 \cdot \frac{e^{\left[\frac{-E}{k}\cdot\left(\frac{1}{T}-\frac{1}{T_{\mathrm{ref}}}\right)\right]}}{1+\frac{E}{E_{\mathrm{D}} - E} \cdot e^{\left[\frac{E_{\mathrm{D}}}{k}\cdot\left(\frac{1}{T_{\mathrm{pk}}}-\frac{1}{T}\right)\right]}}.$$

#### S3.2 Non-positive growth rate values and the Sharpe-Schoolfield model

The Sharpe-Schoolfield model (Eq. (1) in the main text) has an exponential (Arrhenius) term in its numerator and, as a result, the model cannot predict non-positive trait values. Thus, before fitting the model to data, we removed any growth rate measurements $\leq 0$ s$^{-1}$. The only model parameter whose estimate will not be biologically realistic in such a case is $B_0$ (the trait performance at a low reference temperature), which can only take positive values. To assess the impact of this issue on our results, we examined the number of non-positive growth rate measurements at the rise of the TPC (which is partly controlled by $B_0$). Across all species in our final TPC dataset, very few species were exclusively represented by TPCs with negative growth rate values at the rising part (less than 2%). The corresponding number for TPCs that had zero values at the rise was 19%.

For the latter group of TPCs, we also calculated the ratio of trait performance at $T_{\text{ref}} = 0°$C to the maximum trait performance ($B_{\text{pk}}$). The numerator of the ratio (and the ratio itself) should be 0, as in these TPCs, there was at least one zero growth rate measurement at a temperature above 0°C. Indeed, as shown in Fig. S3, the ratio is very close to zero in most cases. This indicates that $B_0$ estimates are, in general, only slight overestimates of true growth rate values and that they do not systematically bias the results of our study.

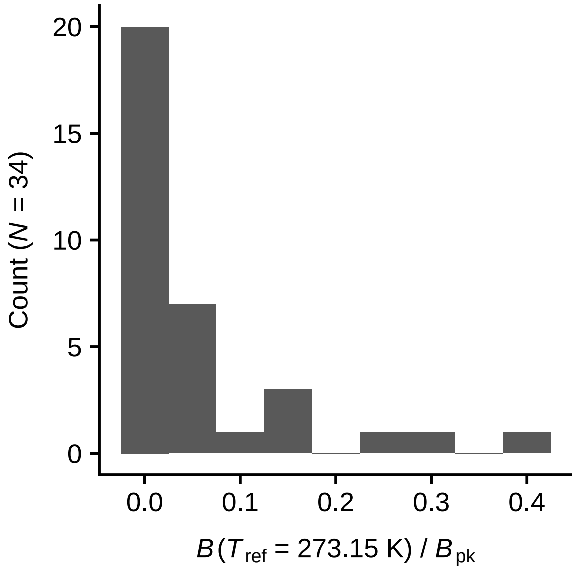

Figure S3: The ratio of $B(T_{\text{ref}})$ to $B_{\text{pk}}$ for experimentally-determined TPCs with a growth rate value of 0 $s^{-1}$ at the rising part. Most values are close to the true ratio of 0.

#### S3.3 Fitting the Sharpe-Schoolfield model to growth rate data

To facilitate nonlinear least squares fitting, we set bounds to three parameters of the model (Table S1). Also, during the optimization process, when the value of $E$ was greater than $E_\mathrm{D}$, we forced the objective function to return $10^{10}$ instead of the expected trait value. This strict penalization forced the nonlinear least squares optimizer to accept only values of $E < E_\mathrm{D}$, in line with experimentally observed thermal performance curves. Additionally, we enforced extremely strict convergence criteria for the optimization, by setting the tolerance parameters xtol and ftol to the value of $10^{-12}$, and the maximum number of iterations to 100,000.

| **Parameter** | **Lower bound** | **Upper bound** |
|---|---|---|
| $E$ | $10^{-5}$ | 10 |
| $E_\mathrm{D}$ | $10^{-5}$ | 30 |
| $T_\mathrm{pk}$ | 263.15 | 423.15 |

Table S1: Parameter bounds set for nonlinear least squares fitting.

We fitted the model to each species or strain whose growth rate had been measured at a minimum of five different temperatures. After fitting the model, we filtered the fits to ensure only reliable and useful parameter estimates – within the context of this study – were retained. To this end, we first rejected i) fits with $R^2 < 0.5$, ii) $E$ estimates that were unrealistically high (i.e., greater than 4 eV), iii) fits where $T_\mathrm{pk} < T_\mathrm{ref}$ (in the case of $T_\mathrm{ref} = 10°\mathrm{C}$), iv) parameter estimates from species that were not in our phylogeny, and v) parameter estimates from species whose isolation location was not available. Next, we ensured that there were enough data points at each area of the thermal performance curve for parameters to be accurately estimated. More precisely, we rejected $B_0$ and $E$ estimates if there were fewer than four data points below $T_\mathrm{pk}$. $T_\mathrm{pk}$ and $B_\mathrm{pk}$ estimates were accepted if there were at least two data points both before and after $T_\mathrm{pk}$. $E_\mathrm{D}$ values were rejected if there were fewer than four data points after $T_\mathrm{pk}$. Finally, to accept $W_\mathrm{op}$ estimates, we required four data points before $T_\mathrm{pk}$ and two after it.

The distributions of the accepted parameter estimates are shown in Figures S4 ($T_\mathrm{ref} = 0°\mathrm{C}$) and S5 ($T_\mathrm{ref} = 10°\mathrm{C}$).

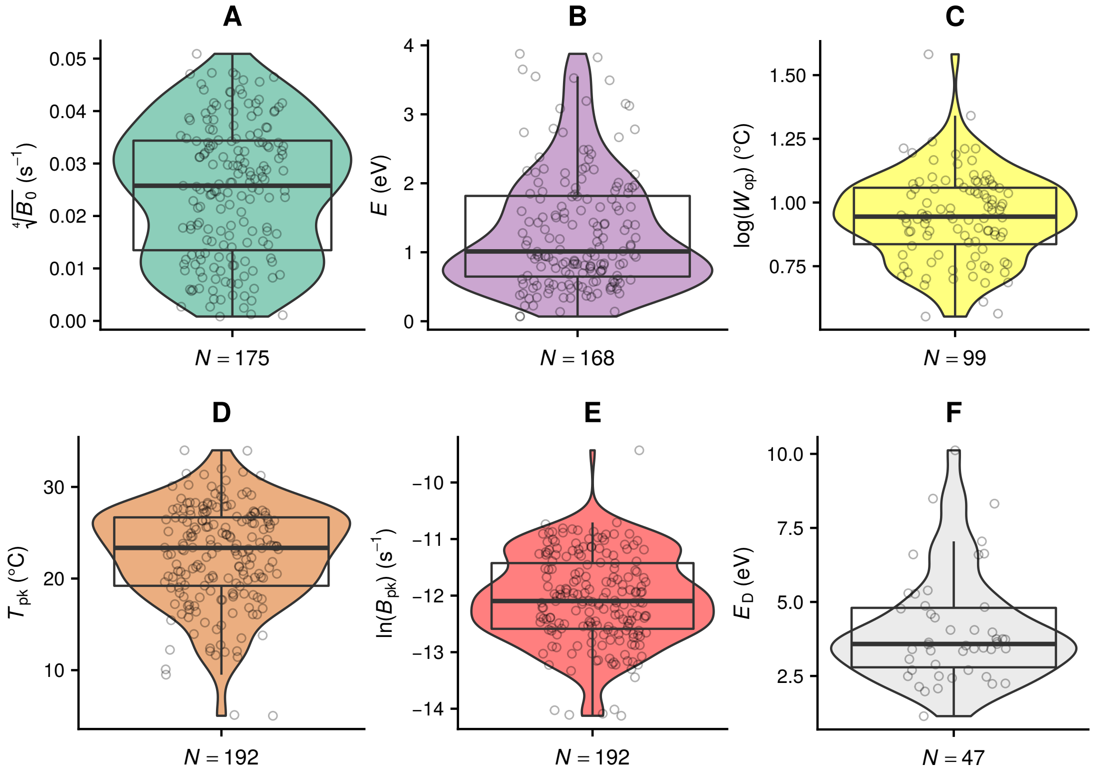

Figure S4: Estimates obtained with a $T_{\mathrm{ref}}$ value of 0°C, for phytoplankton species/strains available in our phylogeny and with known isolation locations.

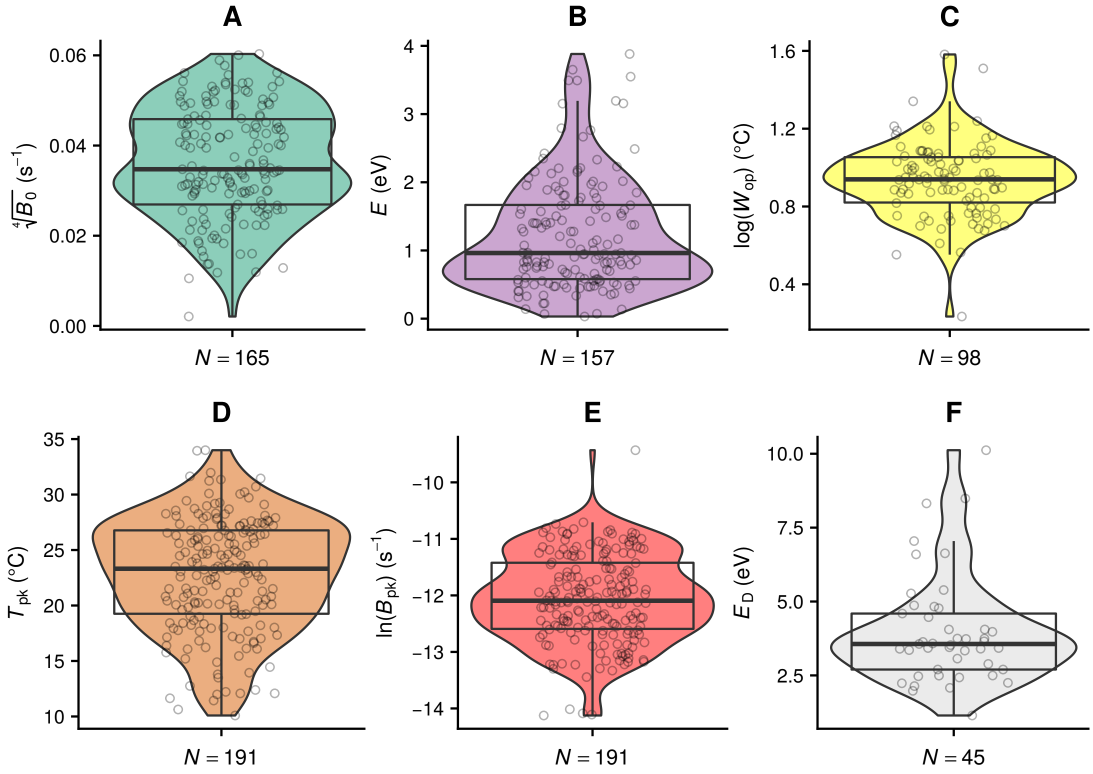

Figure S5: Estimates obtained with a $T_{\mathrm{ref}}$ value of 10°C, and excluding fits with $T_{\mathrm{pk}} < 10$°C, for phytoplankton species/strains available in our phylogeny and with known isolation locations.

### S4 Phylogeny reconstruction diagnostics and tree comparisons

The next two subsections show the outputs of the diagnostics that we ran to verify that the four independent ExaBayes runs (with or without the addition of extra species) had converged.

#### S4.1 Convergence diagnostics for ExaBayes (only required species)

The following diagnostic tests were executed with the AWTY web server (Nylander et al., 2007) and the rwty R package (Warren et al., 2017).

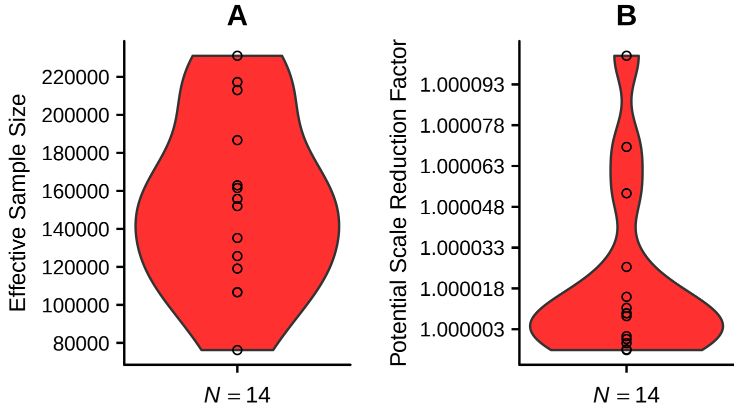

Figure S6: Violin plots of Effective Sample Size (ESS) and Potential Scale Reduction Factor (PSRF) values for the parameters of the evolutionary model across all ExaBayes runs. Convergence and parameter sampling are deemed sufficient if ESS > 200 and PSRF < 1.1.

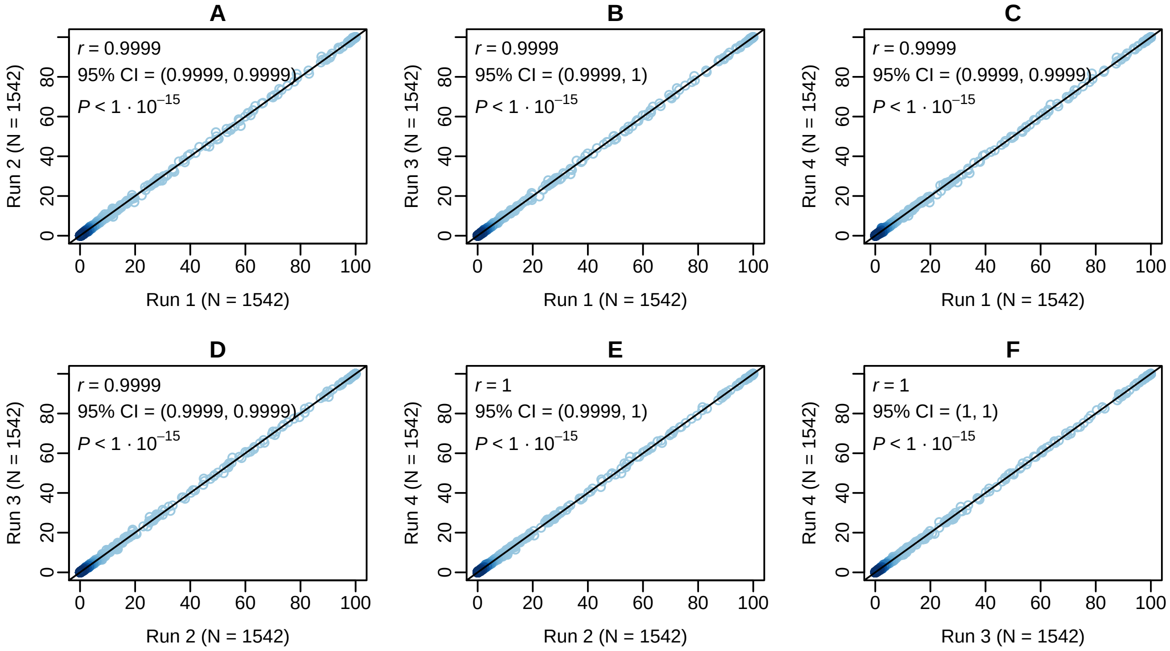

Figure S7: Pairwise comparisons of split frequencies among the four ExaBayes runs, indicating excellent convergence.

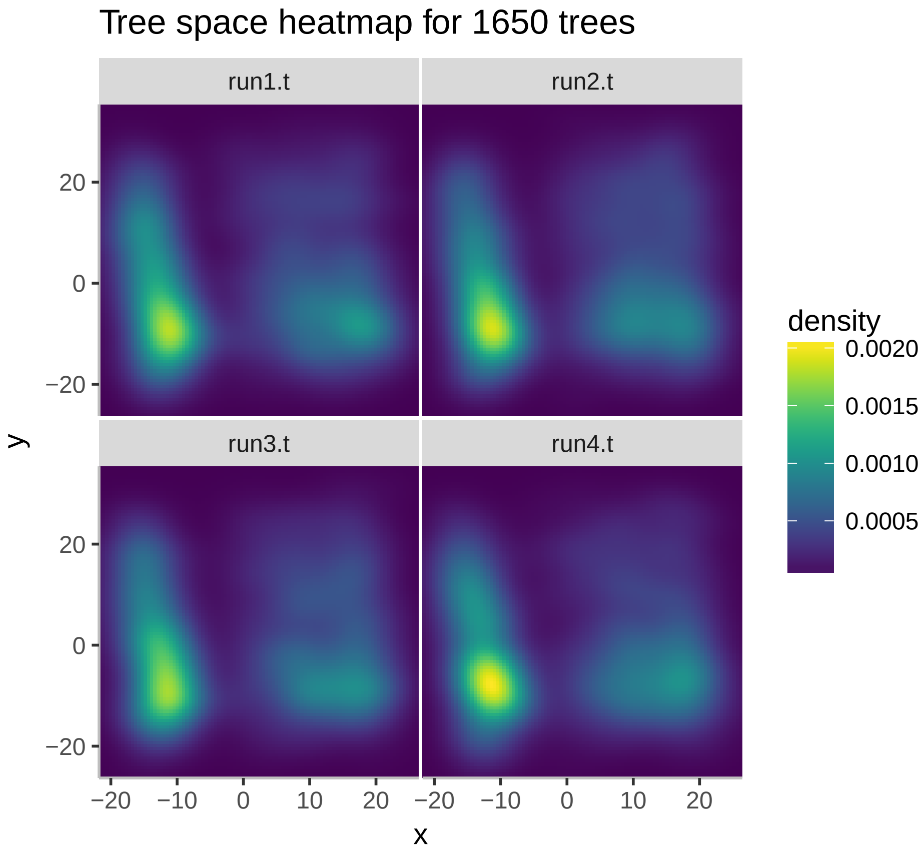

Figure S8: Projection of the sampled tree topologies in two dimensions for each ExaBayes run. As the four panels are highly similar, the runs appear to have adequately converged.

#### S4.2 Convergence diagnostics for ExaBayes (with extra species)

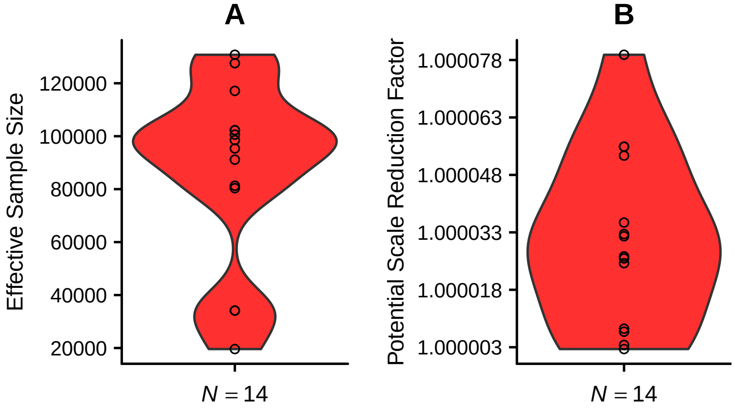

Figure S9: Violin plots of Effective Sample Size (ESS) and Potential Scale Reduction Factor (PSRF) values for the parameters of the evolutionary model across all ExaBayes runs. Convergence and parameter sampling are deemed sufficient if ESS $>$ 200 and PSRF $<$ 1.1.

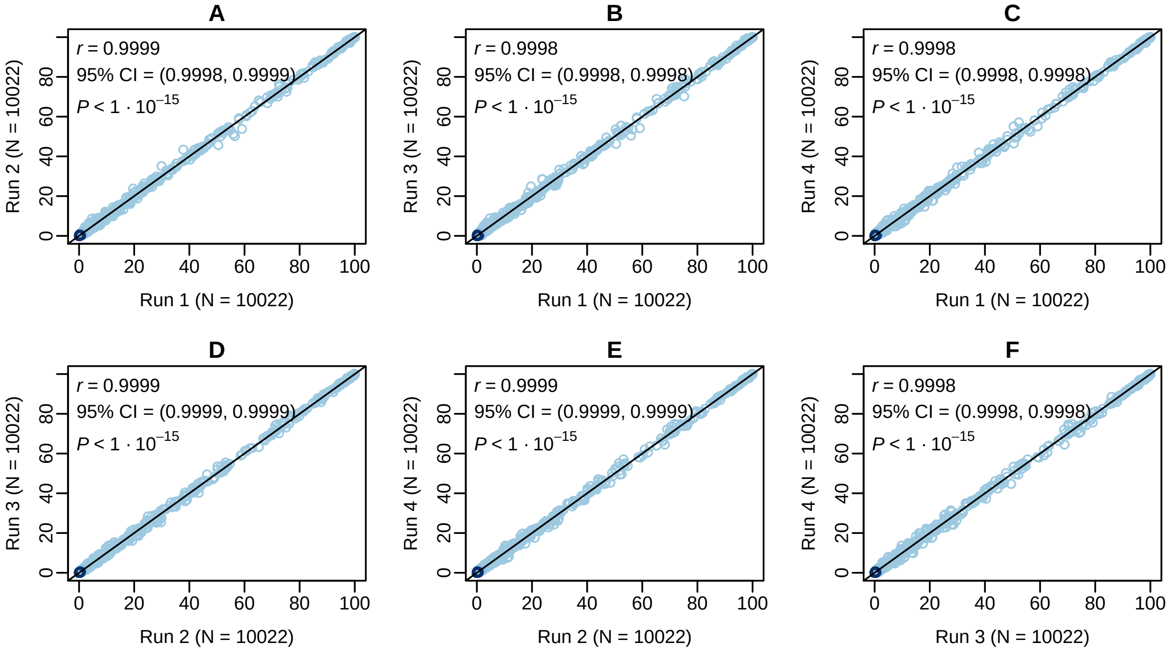

Figure S10: Pairwise comparisons of split frequencies among the four ExaBayes runs, indicating excellent convergence.

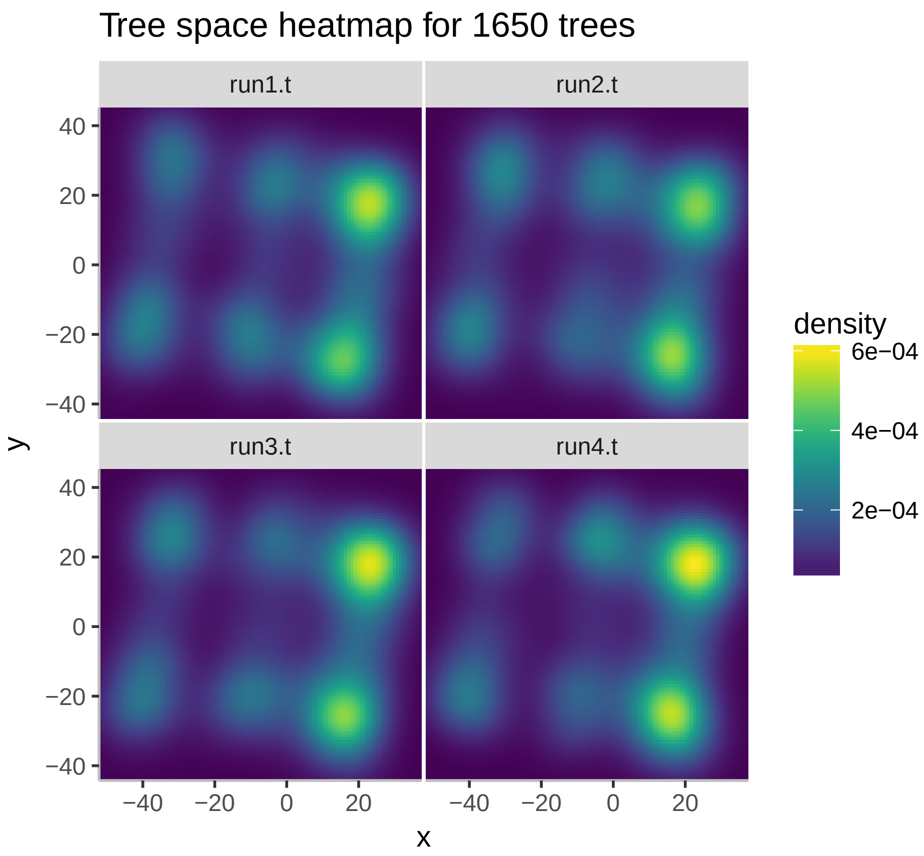

Figure S11: Projection of the sampled tree topologies in two dimensions for each ExaBayes run. As the four panels are highly similar, the runs appear to have adequately converged.

#### S4.3 Comparison of resulting phylogenetic topologies

To understand whether the addition of extra species improved the tree topology by bringing it closer to our knowledge of the tree of life, we compared all resulting topologies to that of the Open Tree of Life (v. 4; Hinchliff et al. 2015). To this end, we first rooted all trees by setting the prokaryotic Cyanophyta species as an outgroup. We then pruned the trees down to a common subset of species. The Open Tree of Life topology was added in two ways: i) including polytomies (i.e., as originally obtained), and ii) with polytomies resolved by RAxML using our alignment of 16S/18S rRNA sequences. Distances between tree topologies were calculated according to the Matching Cluster metric of Bogdanowicz and Giaro (2013), as implemented in TreeCmp (Bogdanowicz et al., 2012). The topologies that were found closest to the Open Tree of Life were those constructed by ExaBayes and RAxML using the sequence alignment with extra species (Fig. S12).

We then compared the log-likelihood of the ExaBayes and RAxML topologies by evaluating them with RAxML under the GTR+Γ model, which had been used for tree reconstruction. As the RAxML topology had a higher log-likelihood (-109356.08) than that of ExaBayes (-109479.86), we used the RAxML tree for the analyses of this study (Fig. S13).

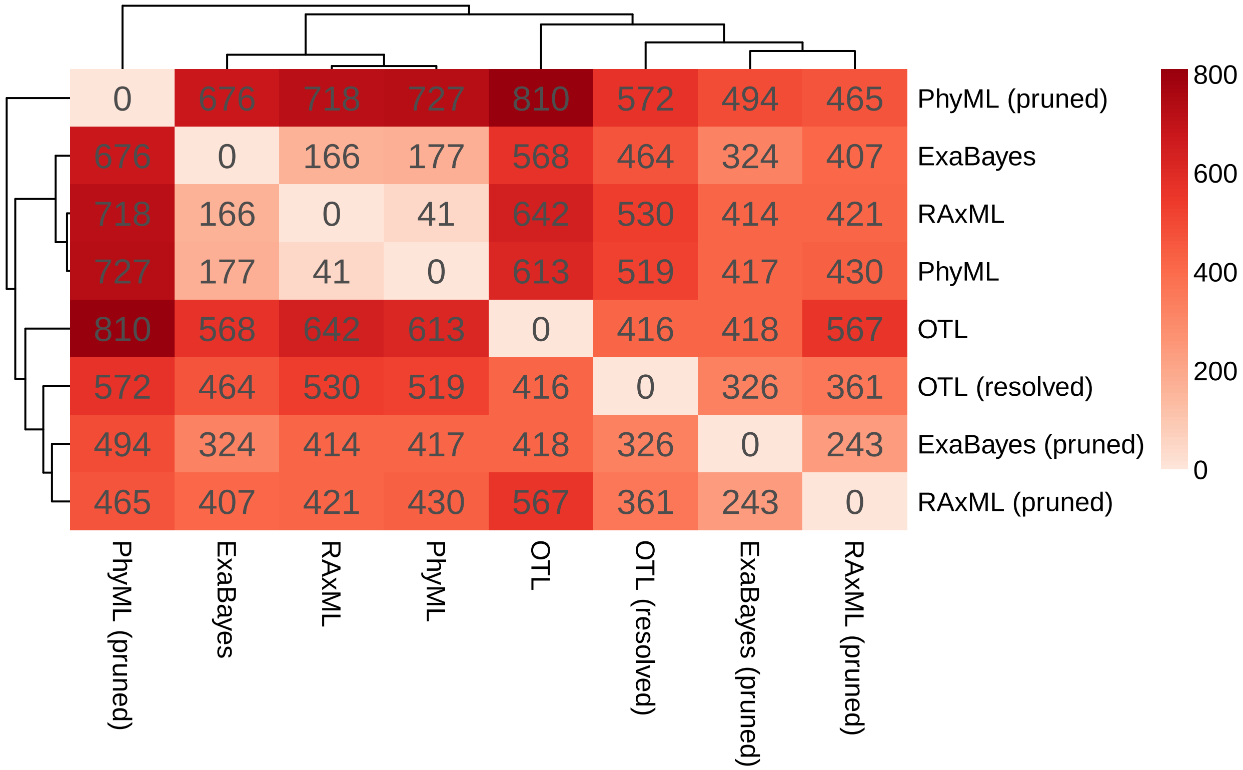

Figure S12: Heat map of Matching Cluster distances between all produced tree topologies and that of the Open Tree of Life (v. 4). Trees marked as "pruned" here are those that included extra species (e.g., macroalgae, land plants). The addition of more species appears to improve the quality of the resulting phylogenetic tree in two of the three programs that we used.

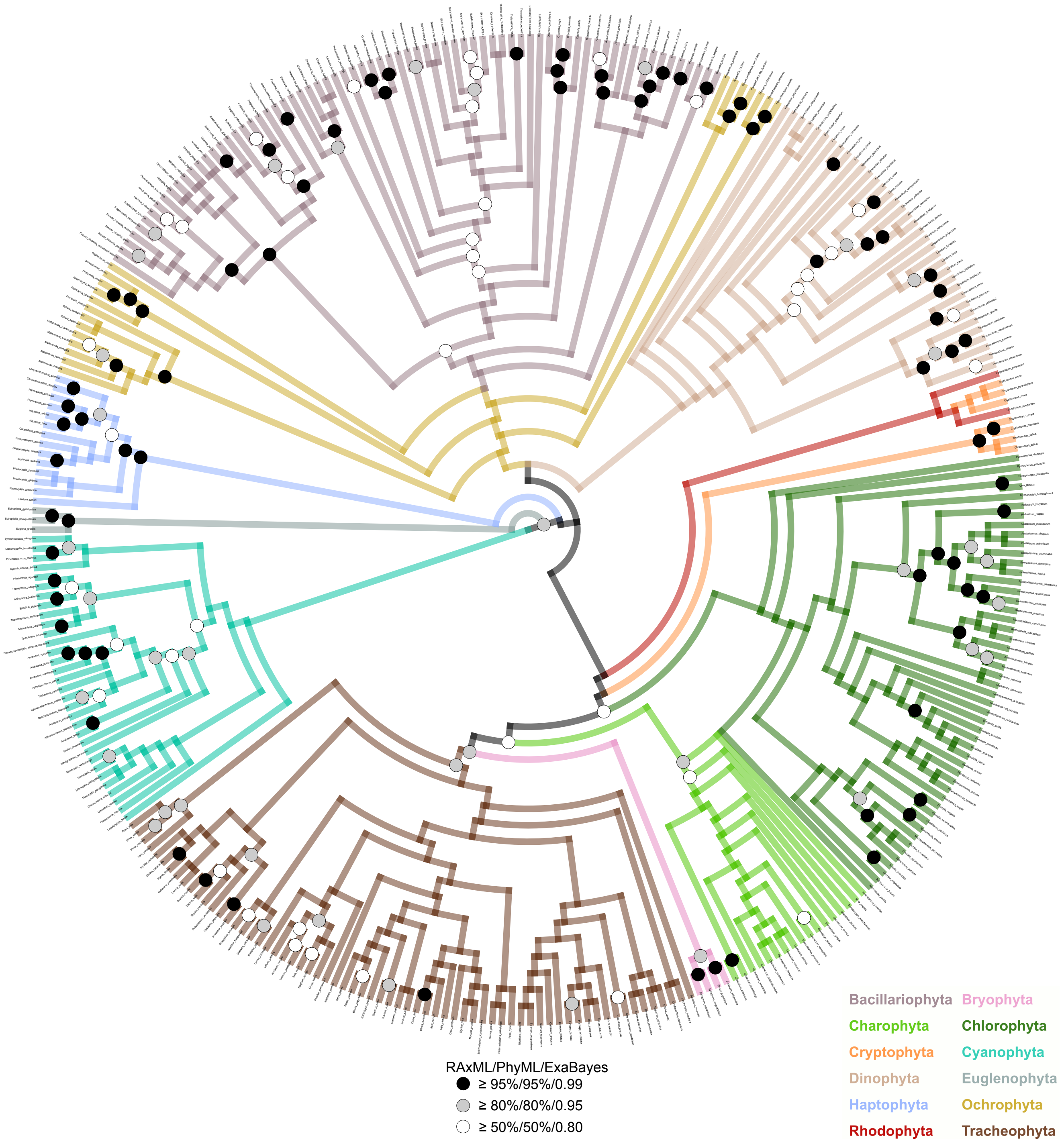

Figure S13: Statistical support – in terms of bootstrap values or posterior probabilities – for the nodes of the extended RAxML topology, according to RAxML, PhyML, and ExaBayes. Nodes with black or grey circles, in particular, are considered robust. Note that, as shown in Fig. S12, the RAxML and ExaBayes topologies are quite close to each other and to the OTL, whereas the PhyML topology is distinctly different. Therefore, the statistical support of many nodes in this figure is probably an underestimate of what we would obtain if we were to exclude the PhyML tree from the comparison.

#### S4.4 Convergence diagnostics for the relative time-calibration (DPP-Div)

As previously, we performed a diagnostic test to verify that the five independent DPPDiv runs had converged on statistically indistinguishable posterior distributions.

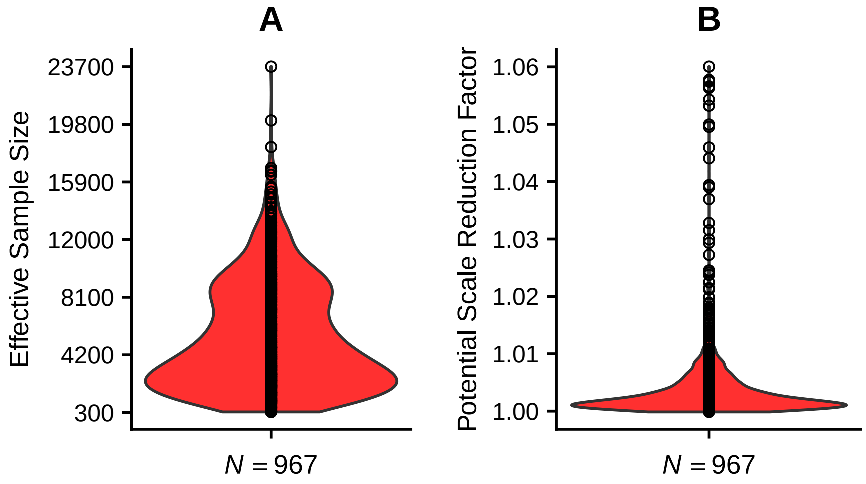

Figure S14: Violin plots of Effective Sample Size (ESS) and Potential Scale Reduction Factor (PSRF) values for the parameters of the relative time-calibration model across the five DPPDiv runs. Convergence and parameter sampling are deemed sufficient if ESS > 200 and PSRF < 1.1.

### S5 Multi-response Markov chain Monte Carlo generalised linear mixed models

#### S5.1 Using the entire dataset

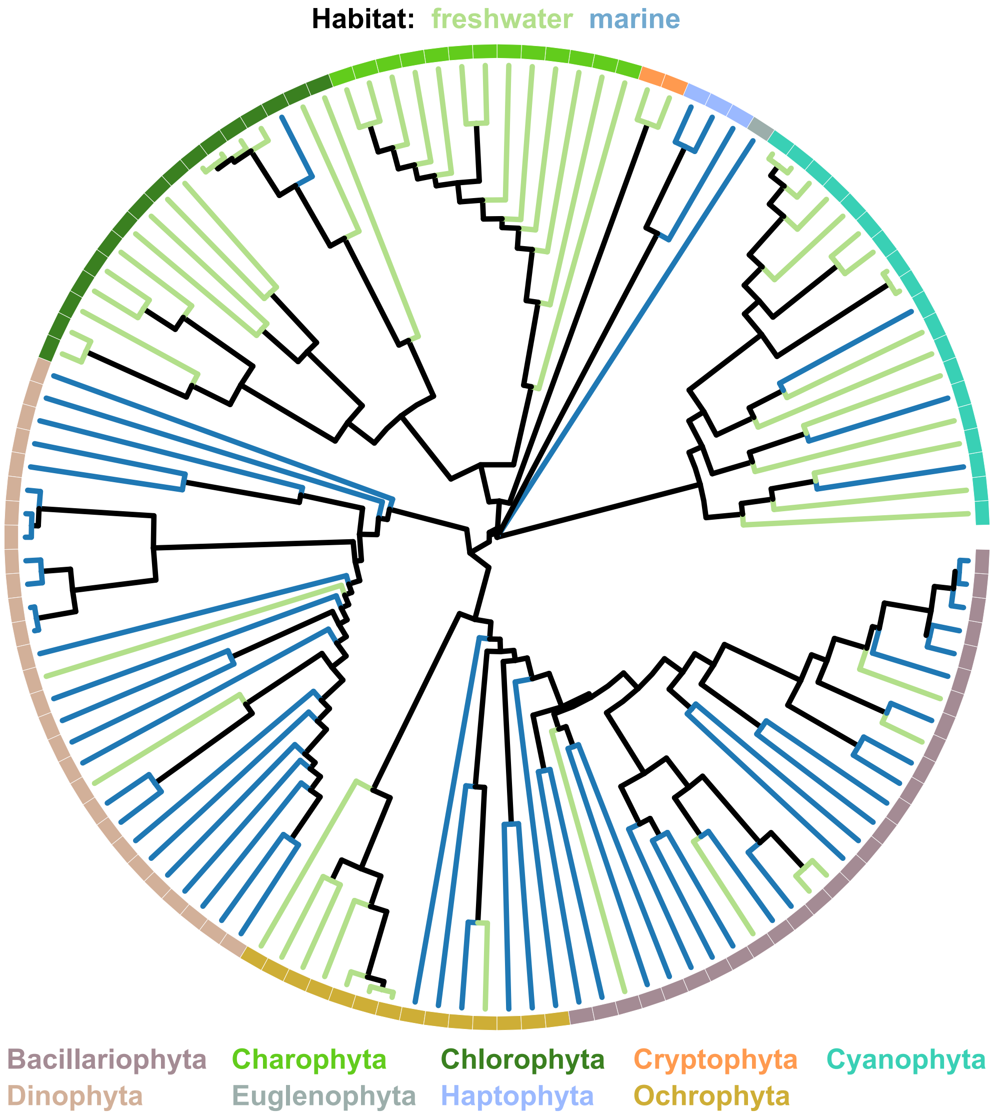

Figure S15: The phylogeny of phytoplankton species in this study, with branches coloured by habitat. The arcs of the circle around the tree denote the various phytoplankton Phyla. This figure was generated using the ggtree R package (Yu et al., 2017).

To assess the extent of phylogenetic sampling bias, we checked whether phylogenetic distance correlates with geographical distance by performing a Mantel test (Mantel, 1967) with 9,999 permutations. In particular, we used the implementation of the Mantel test in the R package ade4 (Dray and Dufour, 2007) to estimate the correlation between the phylogenetic distance matrix and the matrix of geographical distances between species/strains. The latter was calculated using Vincenty's inverse formula (Vincenty, 1975). The correlation between the two matrices was almost nonexistent ($r = 0.0423$, $p = 0.0011$).

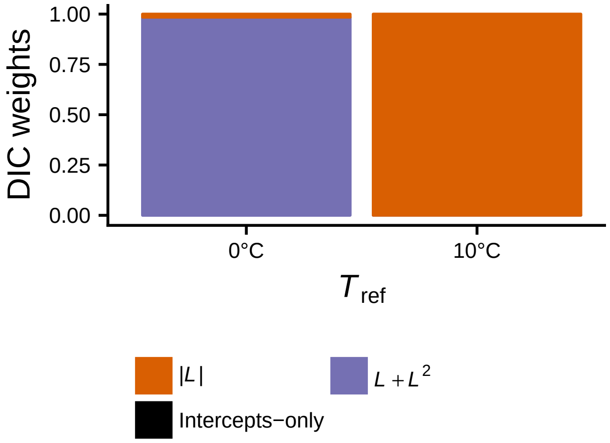

Figure S16: DIC weights of models fitted to the entire dataset of phytoplankton TPC parameters. Models that included the habitat (marine vs freshwater) of the species as a fixed effect are not shown here, as the 95% HPD intervals of the habitat coefficients always contained zero. The best-fitting model had a second-order polynomial in latitude (when $T_{\mathrm{ref}}$ was set to 0°C), or absolute latitude (when $T_{\mathrm{ref}}$ was set to 10°C) as fixed effects.

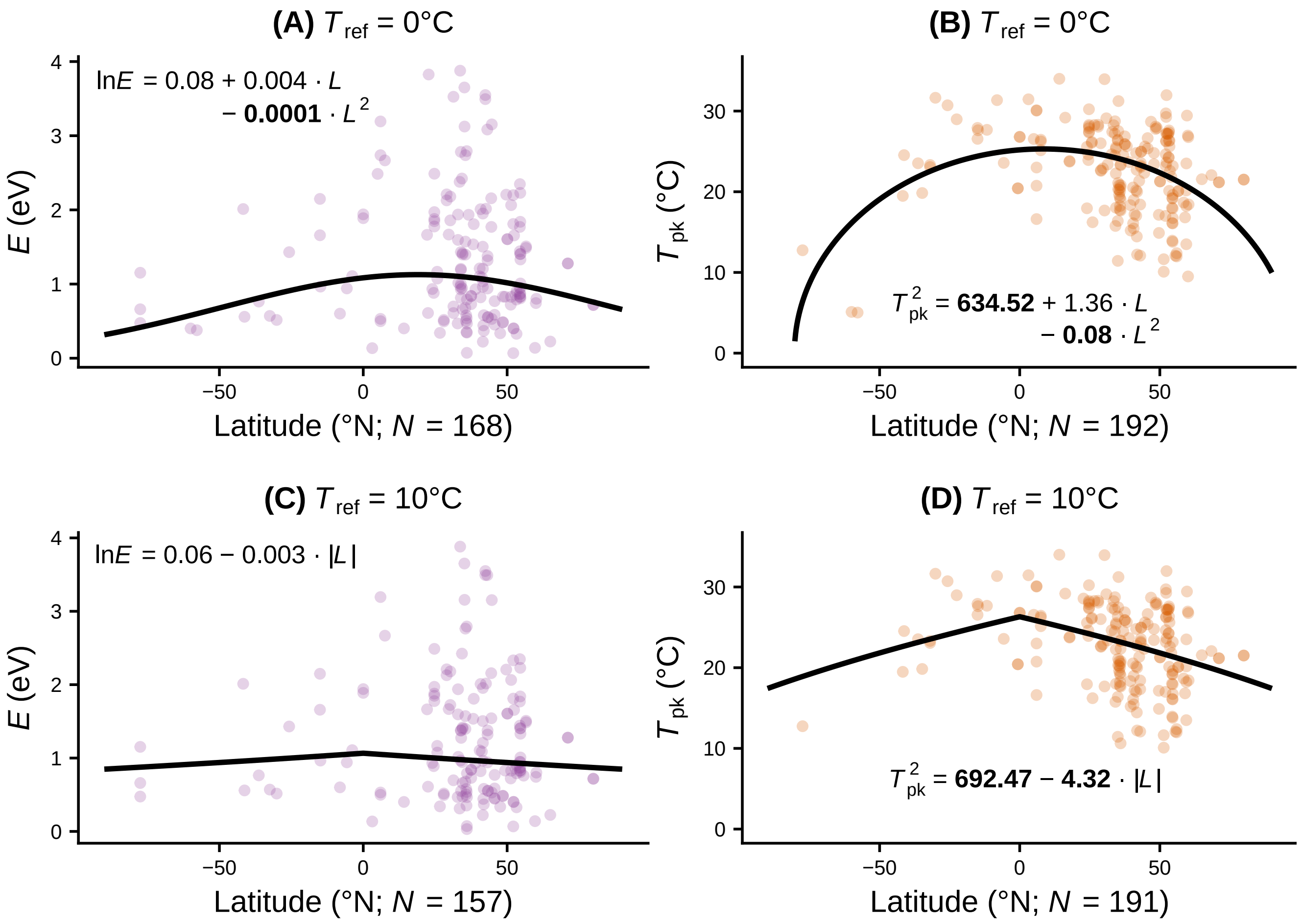

Figure S17: Latitudinal associations of $E$ and $T_{\text{pk}}$. These thermal parameters decrease with absolute latitude, especially if we include species/strains adapted to colder temperatures with $T_{\text{pk}} < 10°\text{C}$. Data points are coloured according to their local density. Coefficients whose 95% Highest Posterior Density interval did not include zero are shown in bold. The lack of the coldest-adapted species changes the best-fitting line from a second-degree polynomial (panels A and B) to a linear relationship with absolute latitude (panels C and D). In panel C, in particular, latitude no longer appears to associate with $E$, as the 95% Highest Posterior Density interval of its coefficient includes zero. These results suggest that, from the equator to higher latitudes, species adapt by both decreasing their $T_{\text{pk}}$ and increasingly becoming thermal generalists.

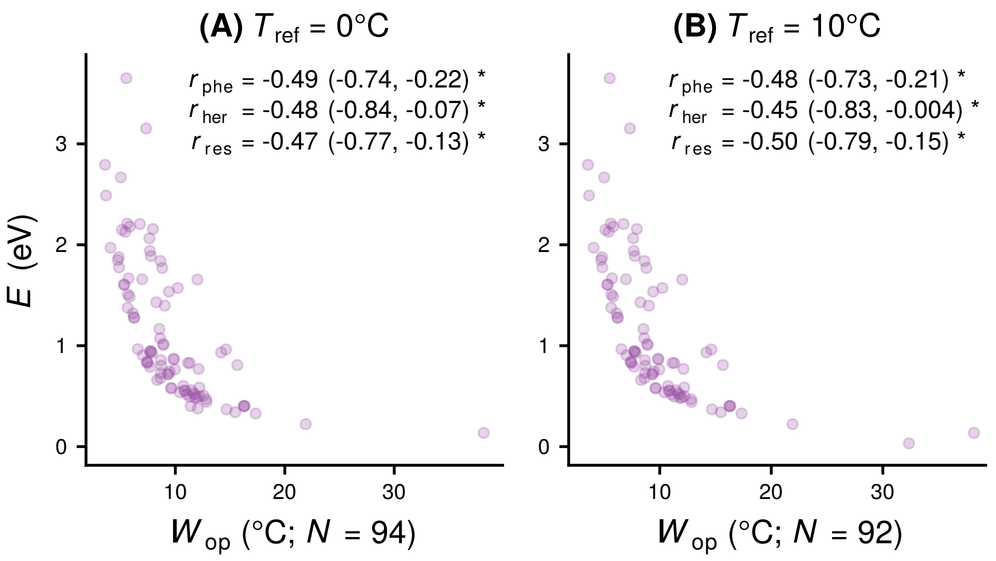

Figure S18: $E$ versus $W_{op}$ using a $T_{ref}$ of 0°C (panel A), or a $T_{ref}$ of 10°C and excluding species/strains with $T_{pk} < 10$°C (panel B). As expected, there is a negative correlation between the two thermal parameters, as TPCs with very steep rising slopes have a narrow operational niche width, and vice versa. The correlation coefficients were estimated between $\ln(E)$ and $\ln(W_{op})$. Performing the analysis with a $T_{ref}$ of 0°C (panel A) or 10°C (panel B) led to almost identical estimates.

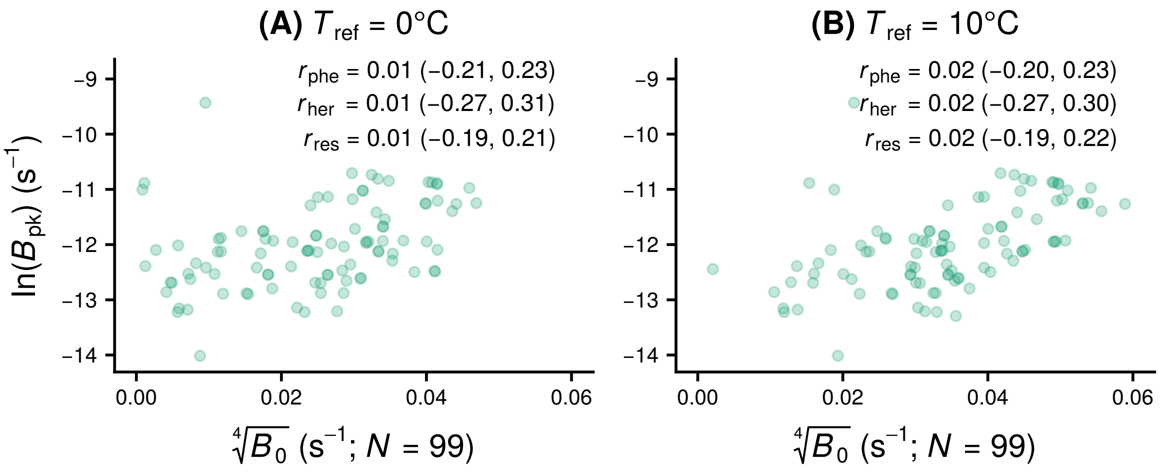

Figure S19: Scatter plots of $B_0$ at $T_{ref}$ = 0°C (panel A) or $T_{ref}$ = 10°C (panel B). The same sample size across the two panels reflects how a change in the $T_{ref}$ value – when fitting the Sharpe-Schoolfield model – can have a minor impact in the value of $R^2$, leading to the acceptance of some fitted curves with $R^2$ values that were previously just below our cutoff of 0.5. While a visual inspection of the plots suggests the presence of a correlation between the two TPC parameters, the correlation becomes non-existent when we control for the phylogenetic distance among species, and the covariance among all TPC parameters. Note that we used different transformations for the two variables in order for their distributions to be as close to normality as possible.

###### S5.1.1 Variance/covariance matrices with $T_{\mathbf{ref}}$ set to 0°C

| | $\sqrt[4]{B_0}$ | $\ln(E)$ | $T_{\mathrm{pk}}^2$ | $\ln(B_{\mathrm{pk}})$ | $\ln(E_{\mathrm{D}})$ | $\ln(W_{\mathrm{op}})$ |
|---|---|---|---|---|---|---|
| $\sqrt[4]{B_0}$ | 0.022<br>(0.015, 0.030) | -0.003<br>(-0.028, 0.022) | -0.415<br>(-10.422, 9.752) | 0.002<br>(-0.034, 0.038) | -0.001<br>(-0.024, 0.022) | 0.002<br>(-0.017, 0.021) |
| $\ln(E)$ | | 0.311<br>(0.102, 0.571) | 21.702<br>(-48.743, 95.562) | -0.029<br>(-0.279, 0.215) | 0.042<br>(-0.142, 0.242) | -0.119<br>(-0.289, 0.018) |
| $T_{\mathrm{pk}}^2$ | | | 54515.880<br>(29333.040, 83886.920) | 75.057<br>(-2.117, 154.759) | 25.155<br>(-50.128, 101.150) | -14.416<br>(-73.133, 42.220) |
| $\ln(B_{\mathrm{pk}})$ | | | | 0.619<br>(0.320, 0.941) | -0.019<br>(-0.268, 0.232) | 0.019<br>(-0.182, 0.223) |
| $\ln(E_{\mathrm{D}})$ | | | | | 0.274<br>(0.085, 0.517) | -0.029<br>(-0.173, 0.109) |
| $\ln(W_{\mathrm{op}})$ | | | | | | 0.188<br>(0.068, 0.343) |

Table S2: Phylogenetically heritable variance/covariance matrix of TPC parameters, as estimated from the intercepts-only model. The first row in each cell stands for the mean value, whereas the second row for the 95% HPD interval.

| | $\sqrt[4]{B_0}$ | $\ln(E)$ | $T_{\mathrm{pk}}^2$ | $\ln(B_{\mathrm{pk}})$ | $\ln(E_{\mathrm{D}})$ | $\ln(W_{\mathrm{op}})$ |
|---|---|---|---|---|---|---|
| $\sqrt[4]{B_0}$ | 0.009<br>(0.007, 0.011) | -0.003<br>(-0.012, 0.007) | 0.001<br>(-0.099, 0.100) | 0.001<br>(-0.009, 0.010) | 0.001<br>(-0.006, 0.007) | 0.001<br>(-0.005, 0.007) |
| $\ln(E)$ | | 0.272<br>(0.174, 0.378) | -0.094<br>(-2.572, 2.356) | -0.014<br>(-0.131, 0.104) | -0.050<br>(-0.152, 0.050) | -0.086<br>(-0.167, -0.008) |
| $T_{\mathrm{pk}}^2$ | | | 44.089<br>(0.067, 158.282) | 0.093<br>(-1.869, 2.106) | 0.023<br>(-1.357, 1.336) | 0.036<br>(-1.216, 1.250) |
| $\ln(B_{\mathrm{pk}})$ | | | | 0.227<br>(0.163, 0.296) | 0.014<br>(-0.076, 0.103) | -0.008<br>(-0.090, 0.072) |
| $\ln(E_{\mathrm{D}})$ | | | | | 0.140<br>(0.064, 0.235) | 0.011<br>(-0.047, 0.072) |
| $\ln(W_{\mathrm{op}})$ | | | | | | 0.110<br>(0.050, 0.183) |

Table S3: Residual variance/covariance matrix of TPC parameters, as estimated from the intercepts-only model. The first row in each cell stands for the mean value, whereas the second row for the 95% HPD interval.

| | $\sqrt[4]{B_0}$ | $\ln(E)$ | $T_{\mathrm{pk}}^2$ | $\ln(B_{\mathrm{pk}})$ | $\ln(E_{\mathrm{D}})$ | $\ln(W_{\mathrm{op}})$ |
|---|---|---|---|---|---|---|
| $\sqrt[4]{B_0}$ | 0.022<br>(0.015, 0.030) | -0.003<br>(-0.029, 0.023) | -0.123<br>(-9.046, 8.602) | 0.002<br>(-0.034, 0.039) | 0.000<br>(-0.024, 0.022) | 0.002<br>(-0.019, 0.021) |
| $\ln(E)$ | | 0.334<br>(0.111, 0.607) | 6.633<br>(-67.701, 81.634) | -0.049<br>(-0.311, 0.212) | 0.018<br>(-0.193, 0.244) | -0.132<br>(-0.313, 0.016) |
| $T_{\mathrm{pk}}^2$ | | | 38585.860<br>(19100.750, 61963.260) | 66.253<br>(-2.078, 137.745) | 16.459<br>(-57.893, 88.398) | -11.019<br>(-68.960, 45.932) |
| $\ln(B_{\mathrm{pk}})$ | | | | 0.626<br>(0.326, 0.956) | 0.000<br>(-0.274, 0.275) | 0.026<br>(-0.189, 0.240) |
| $\ln(E_{\mathrm{D}})$ | | | | | 0.308<br>(0.092, 0.592) | -0.020<br>(-0.184, 0.138) |
| $\ln(W_{\mathrm{op}})$ | | | | | | 0.204<br>(0.072, 0.373) |

Table S4: Phylogenetically heritable variance/covariance matrix of TPC parameters, as estimated from the best fitting model (i.e., with a second order polynomial in latitude as fixed effect).

| | $\sqrt[4]{B_0}$ | $\ln(E)$ | $T_{\mathrm{pk}}^2$ | $\ln(B_{\mathrm{pk}})$ | $\ln(E_{\mathrm{D}})$ | $\ln(W_{\mathrm{op}})$ |
|---|---|---|---|---|---|---|
| $\sqrt[4]{B_0}$ | 0.009<br>(0.007, 0.011) | -0.002<br>(-0.012, 0.007) | 0.002<br>(-0.105, 0.107) | 0.000<br>(-0.009, 0.010) | 0.001<br>(-0.006, 0.007) | 0.001<br>(-0.005, 0.007) |
| $\ln(E)$ | | 0.255<br>(0.161, 0.356) | -0.131<br>(-2.680, 2.342) | -0.005<br>(-0.123, 0.111) | -0.053<br>(-0.153, 0.046) | -0.081<br>(-0.161, -0.007) |
| $T_{\mathrm{pk}}^2$ | | | 56.469<br>(0.069, 188.145) | 0.169<br>(-1.868, 2.536) | 0.041<br>(-1.497, 1.431) | 0.043<br>(-1.302, 1.302) |
| $\ln(B_{\mathrm{pk}})$ | | | | 0.230<br>(0.165, 0.300) | 0.016<br>(-0.077, 0.110) | -0.010<br>(-0.094, 0.073) |
| $\ln(E_{\mathrm{D}})$ | | | | | 0.141<br>(0.060, 0.241) | 0.014<br>(-0.045, 0.076) |
| $\ln(W_{\mathrm{op}})$ | | | | | | 0.110<br>(0.049, 0.182) |

Table S5: Residual variance/covariance matrix of TPC parameters, as estimated from the best fitting model (i.e., with a second order polynomial in latitude as fixed effect).

##### S5.1.2 Variance/covariance matrices with $T_{\mathbf{ref}}$ set to 10°C

| | $\sqrt[4]{B_0}$ | $\ln(E)$ | $T_{\mathrm{pk}}^2$ | $\ln(B_{\mathrm{pk}})$ | $\ln(E_{\mathrm{D}})$ | $\ln(W_{\mathrm{op}})$ |
|---|---|---|---|---|---|---|
| $\sqrt[4]{B_0}$ | 0.023<br>(0.015, 0.031) | -0.002<br>(-0.028, 0.024) | -0.127<br>(-9.623, 9.479) | 0.002<br>(-0.031, 0.036) | 0.000<br>(-0.024, 0.023) | 0.001<br>(-0.018, 0.021) |
| $\ln(E)$ | | 0.310<br>(0.096, 0.588) | -14.496<br>(-85.719, 57.694) | -0.044<br>(-0.269, 0.179) | 0.016<br>(-0.173, 0.216) | -0.114<br>(-0.291, 0.026) |
| $T_{\mathrm{pk}}^2$ | | | 46154.300<br>(23596.560, 72648.160) | 44.455<br>(-26.466, 117.838) | 28.891<br>(-40.279, 98.378) | -3.217<br>(-59.759, 53.247) |
| $\ln(B_{\mathrm{pk}})$ | | | | 0.536<br>(0.264, 0.835) | 0.023<br>(-0.217, 0.260) | 0.016<br>(-0.168, 0.200) |
| $\ln(E_{\mathrm{D}})$ | | | | | 0.269<br>(0.080, 0.511) | -0.021<br>(-0.166, 0.116) |
| $\ln(W_{\mathrm{op}})$ | | | | | | 0.187<br>(0.064, 0.343) |

Table S6: Phylogenetically heritable variance/covariance matrix of TPC parameters, as estimated from the intercepts-only model.

| | $\sqrt[4]{B_0}$ | $\ln(E)$ | $T_{\mathrm{pk}}^2$ | $\ln(B_{\mathrm{pk}})$ | $\ln(E_{\mathrm{D}})$ | $\ln(W_{\mathrm{op}})$ |
|---|---|---|---|---|---|---|
| $\sqrt[4]{B_0}$ | 0.010<br>(0.007, 0.012) | -0.002<br>(-0.012, 0.009) | 0.001<br>(-0.114, 0.107) | 0.001<br>(-0.009, 0.011) | 0.000<br>(-0.007, 0.008) | 0.001<br>(-0.006, 0.007) |
| $\ln(E)$ | | 0.294<br>(0.187, 0.407) | -0.132<br>(-2.974, 2.582) | -0.022<br>(-0.145, 0.106) | -0.054<br>(-0.168, 0.059) | -0.093<br>(-0.181, -0.008) |
| $T_{\mathrm{pk}}^2$ | | | 52.094<br>(0.065, 179.992) | 0.123<br>(-2.058, 2.232) | 0.021<br>(-1.477, 1.516) | 0.050<br>(-1.302, 1.404) |
| $\ln(B_{\mathrm{pk}})$ | | | | 0.235<br>(0.170, 0.306) | 0.007<br>(-0.082, 0.097) | -0.003<br>(-0.088, 0.083) |
| $\ln(E_{\mathrm{D}})$ | | | | | 0.151<br>(0.067, 0.253) | 0.012<br>(-0.053, 0.077) |
| $\ln(W_{\mathrm{op}})$ | | | | | | 0.114<br>(0.051, 0.190) |

Table S7: Residual variance/covariance matrix of TPC parameters, as estimated from the intercepts-only model.

| | $\sqrt[4]{B_0}$ | $\ln(E)$ | $T_{\mathrm{pk}}^2$ | $\ln(B_{\mathrm{pk}})$ | $\ln(E_{\mathrm{D}})$ | $\ln(W_{\mathrm{op}})$ |
|---|---|---|---|---|---|---|
| $\sqrt[4]{B_0}$ | 0.023<br>(0.015, 0.031) | -0.002<br>(-0.028, 0.024) | 0.092<br>(-8.553, 8.958) | 0.002<br>(-0.031, 0.036) | 0.000<br>(-0.024, 0.025) | 0.001<br>(-0.019, 0.021) |
| $\ln(E)$ | | 0.323<br>(0.097, 0.612) | -19.242<br>(-91.345, 50.098) | -0.037<br>(-0.263, 0.190) | 0.011<br>(-0.204, 0.227) | -0.121<br>(-0.305, 0.026) |
| $T_{\mathrm{pk}}^2$ | | | 37610.300<br>(18344.050, 60589.390) | 46.729<br>(-18.112, 115.761) | 18.825<br>(-56.629, 93.119) | -1.325<br>(-57.137, 56.747) |
| $\ln(B_{\mathrm{pk}})$ | | | | 0.532<br>(0.268, 0.838) | 0.038<br>(-0.213, 0.293) | 0.016<br>(-0.171, 0.206) |
| $\ln(E_{\mathrm{D}})$ | | | | | 0.296<br>(0.086, 0.573) | -0.020<br>(-0.181, 0.130) |
| $\ln(W_{\mathrm{op}})$ | | | | | | 0.196<br>(0.066, 0.360) |

Table S8: Phylogenetically heritable variance/covariance matrix of TPC parameters, as estimated from the best fitting model (i.e., with absolute latitude as fixed effect).

| | $\sqrt[4]{B_0}$ | $\ln(E)$ | $T_{\mathrm{pk}}^2$ | $\ln(B_{\mathrm{pk}})$ | $\ln(E_{\mathrm{D}})$ | $\ln(W_{\mathrm{op}})$ |
|---|---|---|---|---|---|---|
| $\sqrt[4]{B_0}$ | 0.010<br>(0.007, 0.012) | -0.002<br>(-0.012, 0.009) | 0.002<br>(-0.120, 0.122) | 0.001<br>(-0.009, 0.011) | 0.000<br>(-0.007, 0.008) | 0.001<br>(-0.006, 0.007) |
| $\ln(E)$ | | 0.294<br>(0.185, 0.408) | -0.199<br>(-3.237, 2.837) | -0.027<br>(-0.149, 0.100) | -0.055<br>(-0.173, 0.058) | -0.094<br>(-0.184, -0.009) |
| $T_{\mathrm{pk}}^2$ | | | 68.851<br>(0.066, 216.891) | 0.184<br>(-2.036, 2.654) | 0.032<br>(-1.647, 1.597) | 0.072<br>(-1.378, 1.564) |
| $\ln(B_{\mathrm{pk}})$ | | | | 0.237<br>(0.170, 0.308) | 0.009<br>(-0.083, 0.102) | 0.001<br>(-0.085, 0.087) |
| $\ln(E_{\mathrm{D}})$ | | | | | 0.149<br>(0.063, 0.255) | 0.015<br>(-0.049, 0.084) |
| $\ln(W_{\mathrm{op}})$ | | | | | | 0.115<br>(0.050, 0.191) |

Table S9: Residual variance/covariance matrix of TPC parameters, as estimated from the best fitting model (i.e., with absolute latitude as fixed effect).

#### S5.2 Using the marine subset of the dataset

##### S5.2.1 With $T_{\text{ref}}$ set to 0°C

Table S10 shows the mean DIC value (averaged across two chains per model) of MCMCglmms fitted on the dataset of marine TPC parameters. Note that a model is included in this table only if each of its fixed effects formed a statistical association with at least one parameter of the TPC (i.e., with at least one response variable). This means that a model was accepted if each of its fixed effects had at least one coefficient whose 95% Highest Posterior Density interval did not include zero. Note that, other than the fixed effects shown in this table, all models also included a distinct intercept for each response variable. The model with the lowest DIC is shown in bold.

Table S10: DIC values for models fitted on the marine subset of the dataset, using a $T_{\text{ref}}$ of 0°C.

| **Fixed effects** | **Mean DIC** |
|---|---|
| - | 283.16 |
| $\lvert L_{\text{orig}}\rvert$ | 156.99 |
| $L_{\text{orig}} + L^2_{\text{orig}}$ | 220.16 |
| $\tilde{T}_{\text{orig}}$ | 179.97 |
| $\tilde{T}_{\text{orig}} + \text{IQR}(T_{\text{orig}}) + \text{IQR}(T_{\text{orig}})^2$ | 209.81 |
| $\tilde{T}_{\text{2.5m, 50d}}$ | 174.82 |
| $\lvert\tilde{L}_{\text{2.5m, 50d}}\rvert$ | 165.89 |
| $\tilde{L}_{\text{2.5m, 50d}} + \tilde{L}^2_{\text{2.5m, 50d}}$ | 191.28 |
| $\tilde{T}_{\text{2.5m, 150d}}$ | 164.03 |
| $\lvert\tilde{L}_{\text{2.5m, 150d}}\rvert$ | 167.12 |
| $\tilde{L}_{\text{2.5m, 150d}} + \tilde{L}^2_{\text{2.5m, 150d}}$ | 189.77 |
| $\tilde{T}_{\text{2.5m, 250d}}$ | 159.90 |
| $\text{IQR}(T_{\text{2.5m, 250d}}) + \text{IQR}(T_{\text{2.5m, 250d}})^2$ | 275.78 |
| $\lvert\tilde{L}_{\text{2.5m, 250d}}\rvert$ | 176.21 |
| $\tilde{L}_{\text{2.5m, 250d}} + \tilde{L}^2_{\text{2.5m, 250d}}$ | 194.92 |
| $\tilde{T}_{\text{2.5m, 350d}}$ | 161.07 |
| $\text{IQR}(T_{\text{2.5m, 350d}}) + \text{IQR}(T_{\text{2.5m, 350d}})^2$ | 272.33 |
| $\tilde{T}_{\text{2.5m, 350d}} + \text{IQR}(T_{\text{2.5m, 350d}})$ | 151.81 |
| $\lvert\tilde{L}_{\text{2.5m, 350d}}\rvert$ | 173.60 |
| $\tilde{L}_{\text{2.5m, 350d}} + \tilde{L}^2_{\text{2.5m, 350d}}$ | 205.54 |
| $\tilde{T}_{\text{2.5m, 500d}}$ | 178.07 |
| $\text{IQR}(T_{\text{2.5m, 500d}}) + \text{IQR}(T_{\text{2.5m, 500d}})^2$ | 269.53 |
| $\tilde{T}_{\text{2.5m, 500d}} + \text{IQR}(T_{\text{2.5m, 500d}})$ | 158.44 |
| $\lvert\tilde{L}_{\text{2.5m, 500d}}\rvert$ | 168.06 |
| $\tilde{L}_{\text{2.5m, 500d}} + \tilde{L}^2_{\text{2.5m, 500d}}$ | 200.88 |
| $\text{IQR}(T_{\text{50m, 50d}})$ | 269.99 |
| $\tilde{T}_{\text{50m, 50d}}$ | 112.37 |
| $\tilde{T}_{\text{50m, 50d}} + \text{IQR}(T_{\text{50m, 50d}})$ | 121.33 |
| $\lvert\tilde{L}_{\text{50m, 50d}}\rvert$ | 181.28 |
| $\tilde{L}_{\text{50m, 50d}} + \tilde{L}^2_{\text{50m, 50d}}$ | 183.27 |
| $\tilde{T}_{\text{50m, 150d}}$ | 112.04 |

Table S10 – *Continued from previous page*

| Fixed effects | Mean DIC |
|---|---|
| $\lvert\tilde{L}_{\text{50m, 150d}}\rvert$ | 174.00 |
| $\tilde{L}_{\text{50m, 150d}} + \tilde{L}^2_{\text{50m, 150d}}$ | 193.19 |
| $\boldsymbol{\tilde{T}_{\text{50m, 250d}}}$ | **110.89** |
| $\lvert\tilde{L}_{\text{50m, 250d}}\rvert$ | 171.43 |
| $\tilde{L}_{\text{50m, 250d}} + \tilde{L}^2_{\text{50m, 250d}}$ | 192.37 |
| $\tilde{T}_{\text{50m, 350d}}$ | 116.82 |
| $\lvert\tilde{L}_{\text{50m, 350d}}\rvert$ | 178.80 |
| $\tilde{L}_{\text{50m, 350d}} + \tilde{L}^2_{\text{50m, 350d}}$ | 186.09 |
| $\tilde{T}_{\text{50m, 500d}}$ | 116.01 |
| $\lvert\tilde{L}_{\text{50m, 500d}}\rvert$ | 174.85 |
| $\tilde{L}_{\text{50m, 500d}} + \tilde{L}^2_{\text{50m, 500d}}$ | 189.50 |
| $\tilde{T}_{\text{100m, 50d}}$ | 202.33 |
| $\lvert\tilde{L}_{\text{100m, 50d}}\rvert$ | 176.64 |
| $\tilde{L}_{\text{100m, 50d}} + \tilde{L}^2_{\text{100m, 50d}}$ | 189.64 |
| $\tilde{T}_{\text{100m, 150d}}$ | 195.92 |
| $\lvert\tilde{L}_{\text{100m, 150d}}\rvert$ | 169.88 |
| $\tilde{L}_{\text{100m, 150d}} + \tilde{L}^2_{\text{100m, 150d}}$ | 202.98 |
| $\tilde{T}_{\text{100m, 250d}}$ | 198.63 |
| $\lvert\tilde{L}_{\text{100m, 250d}}\rvert$ | 167.11 |
| $\tilde{L}_{\text{100m, 250d}} + \tilde{L}^2_{\text{100m, 250d}}$ | 188.91 |
| $\tilde{T}_{\text{100m, 350d}}$ | 202.08 |
| $\lvert\tilde{L}_{\text{100m, 350d}}\rvert$ | 172.11 |
| $\lvert\tilde{L}_{\text{100m, 350d}}\rvert + \text{IQR}(L_{\text{100m, 350d}})$ | 178.78 |
| $\tilde{L}_{\text{100m, 350d}} + \tilde{L}^2_{\text{100m, 350d}}$ | 193.64 |
| $\tilde{T}_{\text{100m, 500d}}$ | 189.05 |
| $\lvert\tilde{L}_{\text{100m, 500d}}\rvert$ | 172.06 |
| $\lvert\tilde{L}_{\text{100m, 500d}}\rvert + \text{IQR}(L_{\text{100m, 500d}})$ | 169.22 |
| $\tilde{L}_{\text{100m, 500d}} + \tilde{L}^2_{\text{100m, 500d}}$ | 194.30 |

The fixed effects of the model with the lowest DIC included i) a distinct intercept for each response variable and ii) $\tilde{T}_{\text{50m, 250d}}$, which we estimated using the simulated Lagrangian trajectories. If we exclude models with variables from drifting trajectories, the best-fitting model had $\lvert L_{\text{orig}}\rvert$ – other than the intercepts – as a fixed effect.

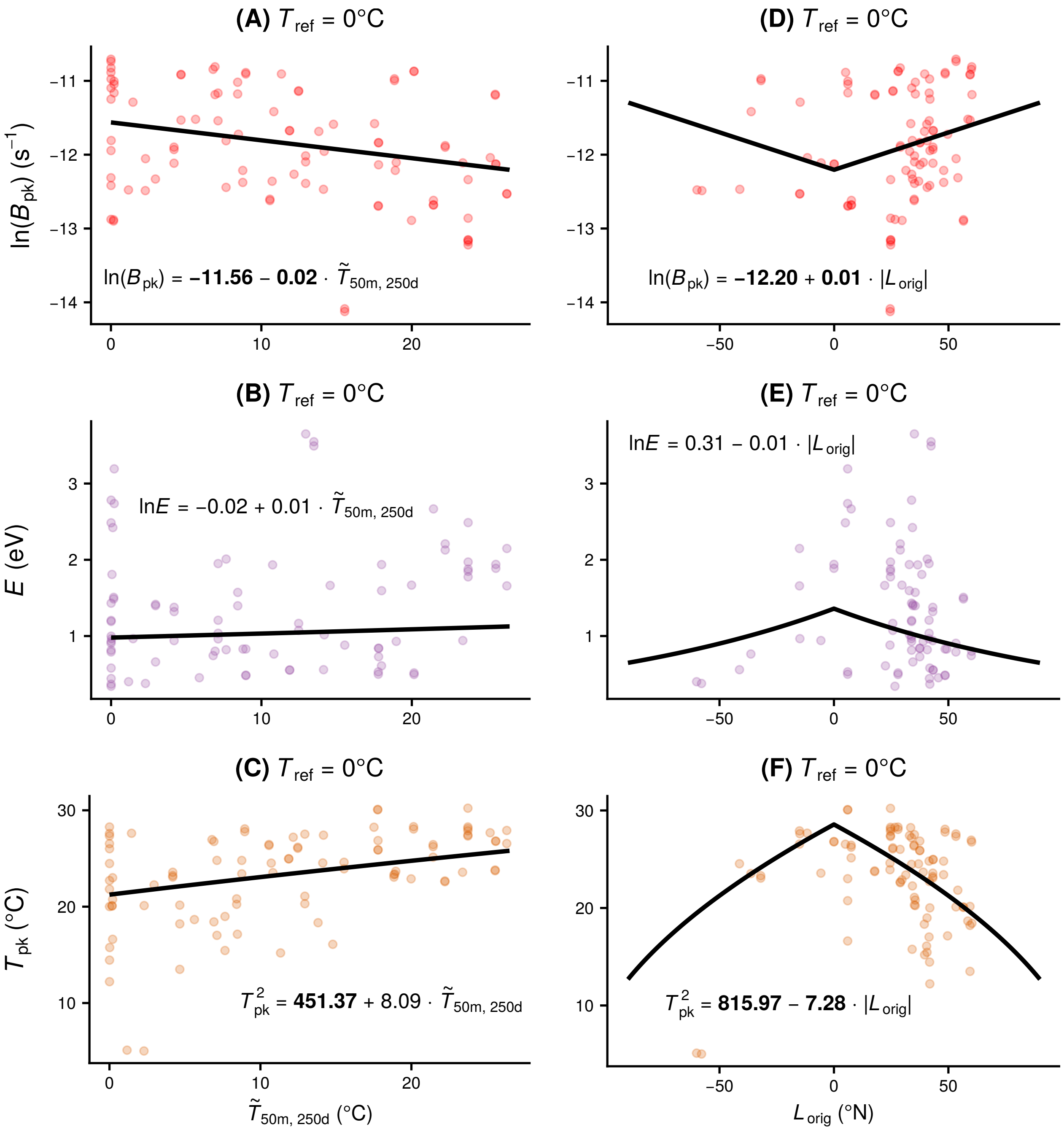

Figure S20: Inferred associations between TPC parameters of marine species/strains and environmental variables. Panels A-C show the relationships of three thermal parameters with the weighted median temperature of estimated drifting trajectories at a depth of 50 meters over 250 days. The MCMCglmm from which the coefficients were obtained for A-C had the lowest DIC among all fitted models. $B_{pk}$ is the only parameter whose $\tilde{T}_{50\text{m, }250\text{d}}$ coefficient had a 95% HPD interval that did not include zero, and is found to decrease with the temperature of the trajectory. This suggests the presence of cold adaptation in marine phytoplankton, with regard to the maximum height of the thermal performance curve. Panels D-F show the inferred relationships of the same parameters using a model with absolute latitude as fixed effect. Here, both $B_{pk}$ and $T_{pk}$ scale with latitude, with the former increasing and the latter decreasing. $E$ does not scale with an environmental variable in any of the two models.

| | $\sqrt[4]{B_0}$ | $\ln(E)$ | $T_{pk}^2$ | $\ln(B_{pk})$ | $\ln(E_D)$ | $\ln(W_{op})$ |
|---|---|---|---|---|---|---|
| $\sqrt[4]{B_0}$ | 0.053<br>(0.027, 0.084) | -0.004<br>(-0.074, 0.063) | -0.450<br>(-24.411, 22.956) | 0.002<br>(-0.068, 0.072) | 0.000<br>(-0.081, 0.080) | 0.002<br>(-0.053, 0.060) |
| $\ln(E)$ | | 0.425<br>(0.116, 0.819) | 21.369<br>(-86.149, 133.125) | -0.060<br>(-0.349, 0.198) | 0.007<br>(-0.367, 0.395) | -0.152<br>(-0.427, 0.071) |
| $T_{pk}^2$ | | | 49591.680<br>(0.111, 87420.630) | 38.675<br>(-52.926, 136.351) | 26.886<br>(-167.099, 230.387) | -16.593<br>(-108.272, 75.629) |
| $\ln(B_{pk})$ | | | | 0.501<br>(0.197, 0.875) | 0.091<br>(-0.327, 0.557) | 0.045<br>(-0.183, 0.287) |
| $\ln(E_D)$ | | | | | 0.574<br>(0.084, 1.430) | -0.003<br>(-0.323, 0.309) |
| $\ln(W_{op})$ | | | | | | 0.285<br>(0.079, 0.574) |

Table S11: Phylogenetically heritable variance/covariance matrix of TPC parameters, as estimated from the best fitting model (i.e., with $\tilde{T}_{50m,\ 250d}$ as fixed effect).

| | $\sqrt[4]{B_0}$ | $\ln(E)$ | $T_{pk}^2$ | $\ln(B_{pk})$ | $\ln(E_D)$ | $\ln(W_{op})$ |
|---|---|---|---|---|---|---|
| $\sqrt[4]{B_0}$ | 0.018<br>(0.012, 0.025) | -0.002<br>(-0.019, 0.015) | 0.012<br>(-0.699, 0.746) | 0.000<br>(-0.022, 0.022) | 0.000<br>(-0.018, 0.019) | 0.001<br>(-0.013, 0.014) |
| $\ln(E)$ | | 0.215<br>(0.113, 0.330) | -0.935<br>(-9.057, 5.939) | 0.000<br>(-0.158, 0.157) | -0.047<br>(-0.188, 0.089) | -0.056<br>(-0.147, 0.030) |
| $T_{pk}^2$ | | | 1244.907<br>(0.060, 5824.667) | 1.910<br>(-6.487, 14.600) | 0.908<br>(-6.949, 10.024) | 0.521<br>(-3.842, 5.712) |
| $\ln(B_{pk})$ | | | | 0.268<br>(0.170, 0.376) | 0.042<br>(-0.144, 0.233) | 0.016<br>(-0.109, 0.138) |
| $\ln(E_D)$ | | | | | 0.249<br>(0.073, 0.487) | 0.013<br>(-0.088, 0.119) |
| $\ln(W_{op})$ | | | | | | 0.127<br>(0.051, 0.223) |

Table S12: Residual variance/covariance matrix of TPC parameters, as estimated from the best fitting model (i.e., with $\tilde{T}_{50m,\ 250d}$ as fixed effect).

| | $\sqrt[4]{B_0}$ | $\ln(E)$ | $T_{\mathrm{pk}}^2$ | $\ln(B_{\mathrm{pk}})$ | $\ln(E_{\mathrm{D}})$ | $\ln(W_{\mathrm{op}})$ |
|---|---|---|---|---|---|---|
| $\sqrt[4]{B_0}$ | 0.053<br>(0.027, 0.084) | -0.004<br>(-0.072, 0.064) | -0.274<br>(-18.690, 17.893) | 0.002<br>(-0.067, 0.073) | 0.001<br>(-0.078, 0.079) | 0.002<br>(-0.055, 0.058) |
| $\ln(E)$ | | 0.431<br>(0.124, 0.812) | 25.421<br>(-57.966, 111.510) | -0.089<br>(-0.371, 0.175) | -0.082<br>(-0.470, 0.268) | -0.160<br>(-0.429, 0.060) |
| $T_{\mathrm{pk}}^2$ | | | 31228.890<br>(8992.526, 66348.630) | 30.729<br>(-41.566, 109.598) | -19.051<br>(-158.646, 120.711) | -19.975<br>(-92.052, 49.631) |
| $\ln(B_{\mathrm{pk}})$ | | | | 0.489<br>(0.179, 0.858) | 0.088<br>(-0.352, 0.564) | 0.068<br>(-0.161, 0.311) |
| $\ln(E_{\mathrm{D}})$ | | | | | 0.562<br>(0.090, 1.336) | 0.050<br>(-0.245, 0.368) |
| $\ln(W_{\mathrm{op}})$ | | | | | | 0.291<br>(0.085, 0.577) |

Table S13: Phylogenetically heritable variance/covariance matrix of TPC parameters, as estimated from the model with $|L_{\mathrm{orig}}|$ as fixed effect.

| | $\sqrt[4]{B_0}$ | $\ln(E)$ | $T_{\mathrm{pk}}^2$ | $\ln(B_{\mathrm{pk}})$ | $\ln(E_{\mathrm{D}})$ | $\ln(W_{\mathrm{op}})$ |
|---|---|---|---|---|---|---|
| $\sqrt[4]{B_0}$ | 0.018<br>(0.012, 0.025) | -0.002<br>(-0.018, 0.015) | 0.003<br>(-0.368, 0.396) | 0.000<br>(-0.022, 0.022) | 0.000<br>(-0.018, 0.019) | 0.001<br>(-0.013, 0.014) |
| $\ln(E)$ | | 0.202<br>(0.109, 0.313) | -0.379<br>(-4.059, 3.256) | 0.003<br>(-0.158, 0.168) | -0.033<br>(-0.172, 0.096) | -0.055<br>(-0.146, 0.026) |
| $T_{\mathrm{pk}}^2$ | | | 450.626<br>(0.061, 765.640) | 0.996<br>(-4.219, 6.134) | 0.234<br>(-3.928, 4.494) | 0.217<br>(-2.294, 2.667) |
| $\ln(B_{\mathrm{pk}})$ | | | | 0.283<br>(0.183, 0.395) | 0.042<br>(-0.147, 0.245) | 0.015<br>(-0.115, 0.142) |
| $\ln(E_{\mathrm{D}})$ | | | | | 0.235<br>(0.066, 0.469) | 0.012<br>(-0.085, 0.120) |
| $\ln(W_{\mathrm{op}})$ | | | | | | 0.127<br>(0.052, 0.223) |

Table S14: Residual variance/covariance matrix of TPC parameters, as estimated from the model with $|L_{\mathrm{orig}}|$ as fixed effect.

##### S5.2.2 With $T_{\text{ref}}$ set to 10°C

As in the previous section, we fitted MCMCglmms to the marine subset of the dataset of TPC parameters, which we obtained by setting $T_{\text{ref}}$ to 10°C.

Table S15: DIC values for models fitted on the marine subset of the dataset, using a $T_{\text{ref}}$ of 10°C.

| **Fixed effects** | **Mean DIC** |
|---|---|
| - | 163.07 |
| $\lvert L_{\text{orig}} \rvert$ | 272.19 |
| $L_{\text{orig}} + L^2_{\text{orig}}$ | 245.28 |
| $\text{IQR}(T_{\text{orig}})$ | 200.76 |
| $\tilde{T}_{\text{orig}}$ | 186.71 |
| $\tilde{T}_{\text{orig}} + \text{IQR}(T_{\text{orig}}) + \text{IQR}(T_{\text{orig}})^2$ | 197.12 |
| $\tilde{T}_{\text{2.5m, 50d}}$ | 236.99 |
| $\lvert \tilde{L}_{\text{2.5m, 50d}} \rvert$ | 239.42 |
| $\tilde{L}_{\text{2.5m, 50d}} + \tilde{L}^2_{\text{2.5m, 50d}}$ | 200.29 |
| $\tilde{T}_{\text{2.5m, 150d}}$ | 185.22 |
| $\lvert \tilde{L}_{\text{2.5m, 150d}} \rvert$ | 248.47 |
| $\tilde{L}_{\text{2.5m, 150d}} + \tilde{L}^2_{\text{2.5m, 150d}}$ | 208.08 |
| $\text{IQR}(T_{\text{2.5m, 250d}})$ | 138.56 |
| $\text{IQR}(T_{\text{2.5m, 250d}}) + \text{IQR}(T_{\text{2.5m, 250d}})^2$ | 155.51 |
| $\tilde{T}_{\text{2.5m, 250d}}$ | 183.60 |
| $\tilde{T}_{\text{2.5m, 250d}} + \text{IQR}(T_{\text{2.5m, 250d}})$ | 451.24 |
| $\lvert \tilde{L}_{\text{2.5m, 250d}} \rvert$ | 237.49 |
| $\tilde{L}_{\text{2.5m, 250d}} + \tilde{L}^2_{\text{2.5m, 250d}}$ | 216.69 |
| $\text{IQR}(T_{\text{2.5m, 350d}})$ | 145.10 |
| $\text{IQR}(T_{\text{2.5m, 350d}}) + \text{IQR}(T_{\text{2.5m, 350d}})^2$ | 148.96 |
| $\tilde{T}_{\text{2.5m, 350d}}$ | 158.83 |
| $\tilde{T}_{\text{2.5m, 350d}} + \text{IQR}(T_{\text{2.5m, 350d}})$ | 413.91 |
| $\lvert \tilde{L}_{\text{2.5m, 350d}} \rvert$ | 251.17 |
| $\tilde{L}_{\text{2.5m, 350d}} + \tilde{L}^2_{\text{2.5m, 350d}}$ | 218.70 |
| $\text{IQR}(T_{\text{2.5m, 500d}})$ | 138.05 |
| $\text{IQR}(T_{\text{2.5m, 500d}}) + \text{IQR}(T_{\text{2.5m, 500d}})^2$ | 144.98 |
| $\tilde{T}_{\text{2.5m, 500d}}$ | 143.74 |
| $\tilde{T}_{\text{2.5m, 500d}} + \text{IQR}(T_{\text{2.5m, 500d}})$ | 359.78 |
| $\lvert \tilde{L}_{\text{2.5m, 500d}} \rvert$ | 246.65 |
| $\tilde{L}_{\text{2.5m, 500d}} + \tilde{L}^2_{\text{2.5m, 500d}}$ | 218.54 |
| $\mathbf{IQR}(\boldsymbol{T}_{\textbf{50m, 50d}})$ | **131.89** |
| $\tilde{T}_{\text{50m, 50d}}$ | 302.47 |
| $\lvert \tilde{L}_{\text{50m, 50d}} \rvert$ | 238.86 |
| $\tilde{L}_{\text{50m, 50d}} + \tilde{L}^2_{\text{50m, 50d}}$ | 216.09 |
| $\tilde{T}_{\text{50m, 150d}}$ | 318.11 |
| $\lvert \tilde{L}_{\text{50m, 150d}} \rvert$ | 244.82 |
| $\tilde{L}_{\text{50m, 150d}} + \tilde{L}^2_{\text{50m, 150d}}$ | 207.26 |

**Table S15** – *Continued from previous page*

| **Fixed effects** | **Mean DIC** |
|---|---|
| $\tilde{T}_{50\text{m, 250d}}$ | 386.06 |
| $\lvert\tilde{L}_{50\text{m, 250d}}\rvert$ | 234.94 |
| $\tilde{L}_{50\text{m, 250d}} + \tilde{L}^2_{50\text{m, 250d}}$ | 212.39 |
| $\tilde{T}_{50\text{m, 350d}}$ | 380.76 |
| $\lvert\tilde{L}_{50\text{m, 350d}}\rvert$ | 260.08 |
| $\tilde{L}_{50\text{m, 350d}} + \tilde{L}^2_{50\text{m, 350d}}$ | 208.43 |
| $\tilde{T}_{50\text{m, 500d}}$ | 396.75 |
| $\lvert\tilde{L}_{50\text{m, 500d}}\rvert$ | 239.78 |
| $\tilde{L}_{50\text{m, 500d}} + \tilde{L}^2_{50\text{m, 500d}}$ | 212.37 |
| $\tilde{T}_{100\text{m, 50d}}$ | 145.56 |
| $\lvert\tilde{L}_{100\text{m, 50d}}\rvert$ | 246.17 |
| $\tilde{L}_{100\text{m, 50d}} + \tilde{L}^2_{100\text{m, 50d}}$ | 215.50 |
| $\tilde{T}_{100\text{m, 150d}}$ | 150.16 |
| $\lvert\tilde{L}_{100\text{m, 150d}}\rvert$ | 245.48 |
| $\tilde{L}_{100\text{m, 150d}} + \tilde{L}^2_{100\text{m, 150d}}$ | 223.21 |
| $\tilde{T}_{100\text{m, 250d}}$ | 153.08 |
| $\lvert\tilde{L}_{100\text{m, 250d}}\rvert$ | 255.41 |
| $\tilde{L}_{100\text{m, 250d}} + \tilde{L}^2_{100\text{m, 250d}}$ | 223.60 |
| $\tilde{T}_{100\text{m, 350d}}$ | 146.88 |
| $\lvert\tilde{L}_{100\text{m, 350d}}\rvert$ | 244.97 |
| $\tilde{L}_{100\text{m, 350d}} + \tilde{L}^2_{100\text{m, 350d}}$ | 217.77 |
| $\tilde{T}_{100\text{m, 500d}}$ | 144.93 |
| $\lvert\tilde{L}_{100\text{m, 500d}}\rvert$ | 244.13 |
| $\tilde{L}_{100\text{m, 500d}} + \tilde{L}^2_{100\text{m, 500d}}$ | 208.25 |

The fixed effects of the best-fitting model overall included i) a distinct intercept for each response variable and ii) $\text{IQR}(T_{50\text{m, 50d}})$, which was obtained from the simulated Lagrangian trajectories. If we were to exclude models with Lagrangian trajectory variables, the model with the lowest DIC was the intercepts-only model.

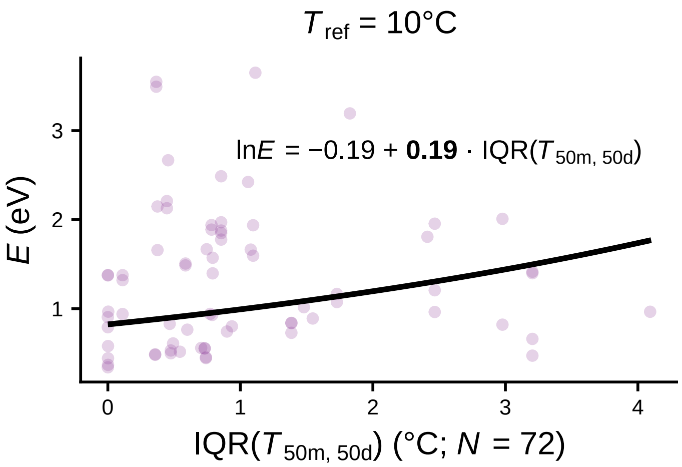

Figure S21: A positive scaling of $E$ with IQR($T_{50m,\ 50d}$) was detected for the marine subset of the dataset and using a $T_{ref}$ of 10°C.

| | $\sqrt[4]{B_0}$ | $\ln(E)$ | $T_{\mathrm{pk}}^2$ | $\ln(B_{\mathrm{pk}})$ | $\ln(E_{\mathrm{D}})$ | $\ln(W_{\mathrm{op}})$ |
|---|---|---|---|---|---|---|
| $\sqrt[4]{B_0}$ | 0.058<br>(0.029, 0.093) | -0.004<br>(-0.081, 0.072) | -0.591<br>(-25.208, 25.056) | 0.004<br>(-0.072, 0.076) | 0.000<br>(-0.080, 0.080) | 0.002<br>(-0.060, 0.065) |
| $\ln(E)$ | | 0.451<br>(0.135, 0.856) | 2.570<br>(-114.448, 121.005) | -0.147<br>(-0.449, 0.111) | 0.004<br>(-0.364, 0.383) | -0.163<br>(-0.448, 0.070) |
| $T_{\mathrm{pk}}^2$ | | | 47998.560<br>(15406.420, 101940.100) | -24.506<br>(-123.608, 73.765) | 13.669<br>(-142.435, 171.837) | -9.701<br>(-108.015, 84.478) |
| $\ln(B_{\mathrm{pk}})$ | | | | 0.472<br>(0.179, 0.834) | -0.014<br>(-0.412, 0.387) | 0.088<br>(-0.132, 0.337) |
| $\ln(E_{\mathrm{D}})$ | | | | | 0.492<br>(0.084, 1.164) | -0.007<br>(-0.314, 0.296) |
| $\ln(W_{\mathrm{op}})$ | | | | | | 0.300<br>(0.082, 0.594) |

Table S16: Phylogenetically heritable variance/covariance matrix of TPC parameters, as estimated from the best fitting model (i.e., with IQR($T_{\mathrm{50m,\ 50d}}$) as fixed effect).

| | $\sqrt[4]{B_0}$ | $\ln(E)$ | $T_{\mathrm{pk}}^2$ | $\ln(B_{\mathrm{pk}})$ | $\ln(E_{\mathrm{D}})$ | $\ln(W_{\mathrm{op}})$ |
|---|---|---|---|---|---|---|
| $\sqrt[4]{B_0}$ | 0.021<br>(0.014, 0.030) | -0.001<br>(-0.021, 0.018) | 0.007<br>(-0.516, 0.574) | 0.001<br>(-0.028, 0.029) | 0.000<br>(-0.021, 0.023) | 0.000<br>(-0.015, 0.016) |
| $\ln(E)$ | | 0.205<br>(0.104, 0.325) | -1.078<br>(-5.989, 3.946) | -0.039<br>(-0.203, 0.123) | -0.055<br>(-0.198, 0.074) | -0.051<br>(-0.142, 0.033) |
| $T_{\mathrm{pk}}^2$ | | | 786.781<br>(0.067, 1209.402) | 0.501<br>(-5.385, 7.145) | 0.514<br>(-4.720, 6.011) | 0.439<br>(-2.933, 3.418) |
| $\ln(B_{\mathrm{pk}})$ | | | | 0.320<br>(0.202, 0.450) | 0.035<br>(-0.196, 0.273) | 0.016<br>(-0.125, 0.159) |
| $\ln(E_{\mathrm{D}})$ | | | | | 0.256<br>(0.070, 0.515) | 0.014<br>(-0.093, 0.125) |
| $\ln(W_{\mathrm{op}})$ | | | | | | 0.128<br>(0.050, 0.224) |

Table S17: Residual variance/covariance matrix of TPC parameters, as estimated from the best fitting model (i.e., with IQR($T_{\mathrm{50m,\ 50d}}$) as fixed effect).

| | $\sqrt[4]{B_0}$ | $\ln(E)$ | $T_{\text{pk}}^2$ | $\ln(B_{\text{pk}})$ | $\ln(E_{\text{D}})$ | $\ln(W_{\text{op}})$ |
|---|---|---|---|---|---|---|
| $\sqrt[4]{B_0}$ | 0.057<br>(0.029, 0.093) | -0.003<br>(-0.081, 0.073) | -0.554<br>(-25.419, 24.524) | 0.003<br>(-0.068, 0.076) | 0.000<br>(-0.078, 0.077) | 0.002<br>(-0.059, 0.062) |
| $\ln(E)$ | | 0.457<br>(0.122, 0.881) | -0.078<br>(-120.612, 115.701) | -0.126<br>(-0.427, 0.137) | -0.009<br>(-0.373, 0.373) | -0.158<br>(-0.446, 0.075) |
| $T_{\text{pk}}^2$ | | | 49565.850<br>(17809.710, 93889.040) | -21.044<br>(-119.439, 76.093) | -6.786<br>(-152.725, 141.140) | -10.779<br>(-104.098, 84.569) |
| $\ln(B_{\text{pk}})$ | | | | 0.455<br>(0.170, 0.815) | -0.054<br>(-0.305, 0.423) | 0.080<br>(-0.138, 0.321) |
| $\ln(E_{\text{D}})$ | | | | | 0.469<br>(0.080, 1.088) | 0.008<br>(-0.283, 0.295) |
| $\ln(W_{\text{op}})$ | | | | | | 0.291<br>(0.080, 0.578) |

Table S18: Phylogenetically heritable variance/covariance matrix of TPC parameters, as estimated from the intercepts-only model.

| | $\sqrt[4]{B_0}$ | $\ln(E)$ | $T_{\text{pk}}^2$ | $\ln(B_{\text{pk}})$ | $\ln(E_{\text{D}})$ | $\ln(W_{\text{op}})$ |
|---|---|---|---|---|---|---|
| $\sqrt[4]{B_0}$ | 0.021<br>(0.014, 0.029) | -0.001<br>(-0.022, 0.019) | 0.002<br>(-0.378, 0.387) | 0.001<br>(-0.028, 0.029) | 0.001<br>(-0.023, 0.024) | 0.000<br>(-0.015, 0.016) |
| $\ln(E)$ | | 0.231<br>(0.119, 0.367) | -0.653<br>(-4.550, 3.190) | -0.052<br>(-0.241, 0.151) | -0.070<br>(-0.238, 0.086) | -0.060<br>(-0.165, 0.032) |
| $T_{\text{pk}}^2$ | | | 412.268<br>(0.065, 528.633) | 0.481<br>(-4.301, 5.433) | 0.371<br>(-3.861, 4.949) | 0.279<br>(-1.980, 2.533) |
| $\ln(B_{\text{pk}})$ | | | | 0.328<br>(0.209, 0.462) | 0.084<br>(-0.147, 0.322) | 0.023<br>(-0.125, 0.170) |
| $\ln(E_{\text{D}})$ | | | | | 0.288<br>(0.085, 0.575) | 0.018<br>(-0.101, 0.137) |
| $\ln(W_{\text{op}})$ | | | | | | 0.130<br>(0.050, 0.230) |

Table S19: Residual variance/covariance matrix of TPC parameters, as estimated from the intercepts-only model.

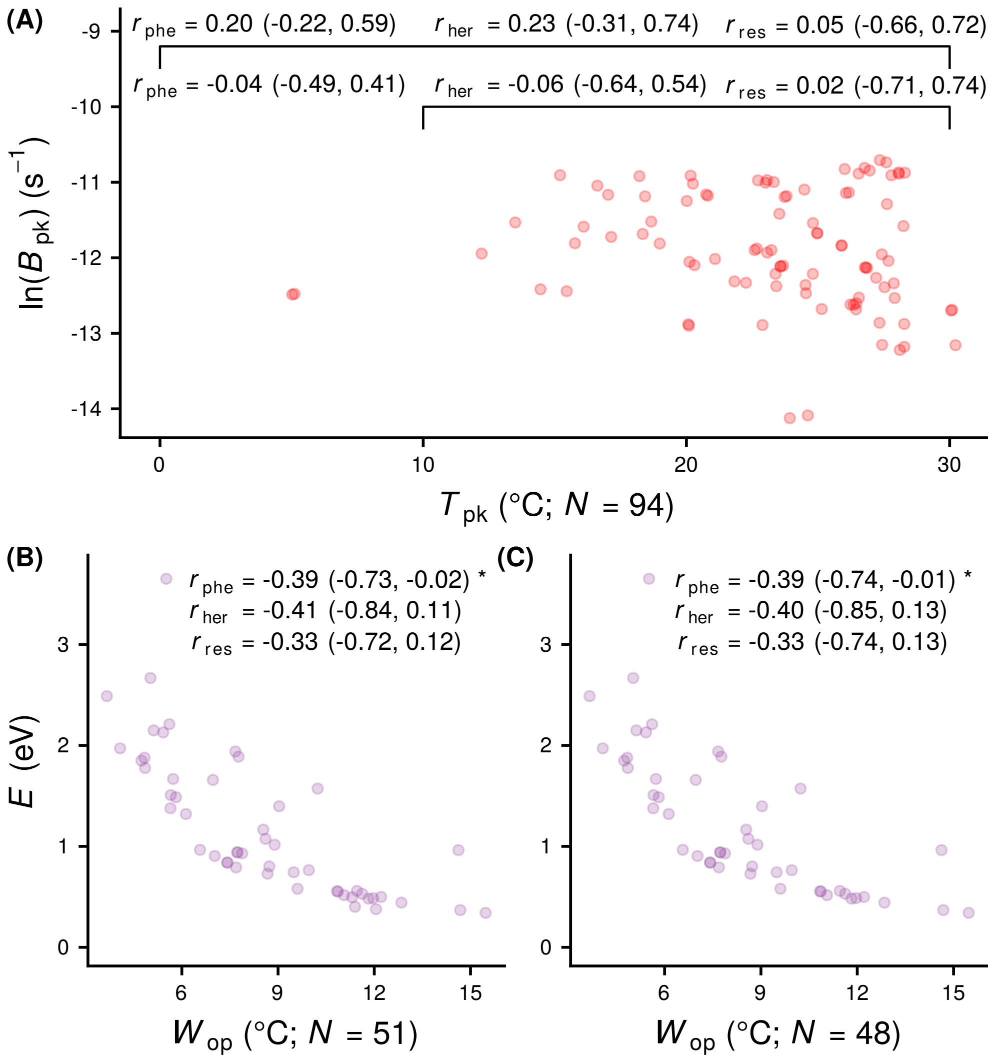

Figure S22: Inferred correlations between TPC parameters of marine species/strains. A: A positive correlation between $T_{\mathrm{pk}}$ and $B_{\mathrm{pk}}$ – as suggested by the "hotter-is-better" hypothesis – could not be detected. This is partly expected from figure S20, as $B_{\mathrm{pk}}$ increases with absolute latitude, whereas $T_{\mathrm{pk}}$ decreases. B-C: A correlation between $\ln(E)$ and $\ln(W_{\mathrm{op}})$ can still be detected regardless of $T_{\mathrm{ref}}$ value (0°C in panel B, 10°C in C), albeit marginally, possibly due to the smaller sample size.

### S6 Scaling of $B_0$ and $B_{\mathrm{pk}}$ with cell volume

To detect any effects of cell volume on growth rate at low temperatures ($B_0$) or on the maximum possible growth rate for each TPC ($B_{\mathrm{pk}}$), we performed the regressions shown in Table S20 using the MCMCglmm R package. The models with the lowest DIC for each response variable are shown in Fig. 5 in the main text.

Table S20: DIC values for regressions of $\sqrt[4]{B_0}$ or $\ln(B_{\mathrm{pk}})$ against the natural logarithm of cell volume.

| **Model structure** | **Response variable** | | | |
|---|---|---|---|---|
| | $T_{\mathrm{ref}} = 0°\mathrm{C}$ | | $T_{\mathrm{ref}} = 10°\mathrm{C}$ | |
| | $\sqrt[4]{\boldsymbol{B_0}}$ | $\mathbf{ln}(\boldsymbol{B_{\mathrm{pk}}})$ | $\sqrt[4]{\boldsymbol{B_0}}$ | $\mathbf{ln}(\boldsymbol{B_{\mathrm{pk}}})$ |
| Species identity as a random effect on the intercept | -473.50 | **316.82** | -424.18 | **314.76** |
| Species identity as a random effect on the intercept, phylogenetic correction | **-518.06** | 328.00 | **-467.62** | 327.02 |
| Species identity as a random effect on the slope | -388.56 | 329.40 | -336.52 | 326.43 |
| Species identity as a random effect on the slope, phylogenetic correction | -389.57 | 328.22 | -337.66 | 325.46 |
| Species identity as a random effect on the intercept and the slope | -388.55 | 329.55 | -336.50 | 326.60 |
| Species identity as a random effect on the intercept and the slope, phylogenetic correction | -389.42 | 328.32 | -337.64 | 325.49 |

### S7 List of 16S/18S rRNA sequences used for phylogeny reconstruction

Table S21: Species names and Accession IDs of sequences that were used in this study.

| **Species** | **Accession ID** |
|---|---|
| *Abies alba* | GenBank: DQ371809.1 |
| *Abutilon theophrasti* | GenBank: DQ287985.1 |
| *Acer rubrum* | GenBank: U42494.1 |
| *Acutodesmus obliquus* | GenBank: KC852905.1 |
| *Aldrovanda vesiculosa* | GenBank: AY096114.1 |
| *Alexandrium catenella* | GenBank: AJ535392.1 |
| *Alexandrium fundyense* | GenBank: KF908796.1 |
| *Alexandrium minutum* | GenBank: U27499.1 |
| *Alexandrium monilatum* | GenBank: AY883005.1 |
| *Alexandrium ostenfeldii* | GenBank: U27500.1 |
| *Alexandrium tamarense* | GenBank: AJ415510.1 |
| *Alexandrium tamiyavanichi* | GenBank: AB088325.1 |
| *Amphidinium klebsii* | GenBank: EU046335.1 |
| *Amphiprora paludosa* | GenBank: AY485468.1 |
| *Anabaena bergii* | GenBank: AF160256.1 |
| *Anabaena cylindrica* | GenBank: HF678516.1 |
| *Anabaena macrospora* | GenBank: AJ293115.1 |
| *Anabaena spiroides* | GenBank: AB271212.1 |
| *Anabaena ucrainica* | GenBank: AB551452.1 |
| *Ankistrodesmus falcatus* var. *tumidus* | GenBank: JQ315498.1 |
| *Apedinella radians* | GenBank: U14384.1 |
| *Aphanizomenon gracile* | GenBank: AJ293127.1 |
| *Aphanizomenon ovalisporum* | GenBank: FM177484.1 |
| *Aplectrum hyemale* | GenBank: U59937.1 |
| *Arbutus unedo* | GenBank: AF206853.1 |
| *Aristotelia serrata* | GenBank: GU476422.1 |
| *Arthrospira fusiformis* | GenBank: AF260510.2 |
| *Asterionella formosa* | GenBank: AM712617.1 |
| *Asterionellopsis glacialis* | GenBank: X77701.1 |
| *Aulacoseira baicalensis* | GenBank: AY121821.1 |
| *Aulacoseira granulata* | GenBank: AB430586.1 |
| *Aulacoseira subarctica* | GenBank: AY121818.1 |
| *Betula papyrifera* | GenBank: L00971.1 |
| *Betula pendula* | GenBank: GU476453.1 |
| *Botryococcus braunii* | GenBank: AB780365.1 |
| *Brassica oleracea* | GenBank: KJ607174.1 |
| *Brassica rapa* | GenBank: LC009534.1 |
| *Bryum argenteum* | GenBank: U18529.1 |
| *Capsicum annuum* | GenBank: EF564281.1 |

**Table S21** – *Continued from previous page*

| Species | Accession ID |
|---|---|
| *Carya glabra* | GenBank: AF206880.1 |
| *Caulerpa serrulata* | GenBank: JQ745683.1 |
| *Ceratium furca* | GenBank: AJ276699.1 |
| *Ceratium furcoides* | GenBank: JQ639757.1 |
| *Ceratium fusus* | GenBank: AF022153.1 |
| *Ceratophyllum demersum* | GenBank: U42517.1 |
| *Chaetoceros calcitrans* | GenBank: JF489975.1 |
| *Chaetoceros debilis* | GenBank: AB847419.1 |
| *Chaetoceros didymus* | GenBank: X85392.2 |
| *Chaetoceros gracilis* | GenBank: AY625895.1 |
| *Chamaebatiaria millefolium* | GenBank: DQ886366.1 |
| *Chatonella marina* | GenBank: AB217627.1 |
| *Chlamydomonas acidophila* | GenBank: AJ852427.1 |
| *Chlamydomonas globosa* | GenBank: AB753039.1 |
| *Chlamydomonas reinhardtii* | GenBank: KF864473.1 |
| *Chlamydomonas subcaudata* | GenBank: AJ781310.1 |
| *Chlorella ellipsoidea* | GenBank: X63520.1 |
| *Chlorella pyrenoidosa* | GenBank: AB240151.1 |
| *Chlorella saccharophila* | GenBank: AB183577.1 |
| *Chlorella sorokiniana* | GenBank: EU402596.1 |
| *Chlorella vulgaris* | GenBank: HQ702325.1 |
| *Chroococcus minutus* | GenBank: GQ375047.1 |
| *Chroomonas salina* | GenBank: GU983864.1 |
| *Chrysanthemum morifolium* | GenBank: KJ870235.1 |
| *Chrysochromulina acantha* | GenBank: AJ246278.1 |
| *Chrysochromulina simplex* | GenBank: AM491021.2 |
| *Cicer arietinum* | GenBank: AJ011011.4 |
| *Citrus aurantium* | GenBank: U38312.1 |
| *Citrus limon* | GenBank: KJ740202.1 |
| *Cladophora glomerata* | GenBank: AB665579.1 |
| *Closterium acerosum* | GenBank: AF352230.1 |
| *Closterium ehrenbergii* | GenBank: AF352228.1 |
| *Coccolithus pelagicus* | GenBank: AJ246261.1 |
| *Cochlodinium polykrikoides* | GenBank: EU418971.1 |
| *Coelastrum astroideum* | GenBank: AF388380.1 |
| *Coelastrum microporum* | GenBank: JQ315527.1 |
| *Coolia monotis* | GenBank: EF492487.1 |
| *Coscinodiscus concinnus* | GenBank: HQ912681.1 |
| *Coscinodiscus granii* | GenBank: AY485495.1 |
| *Coscinodiscus jonesianus* | GenBank: KJ577852.1 |
| *Coscinodiscus wailesii* | GenBank: HQ912668.1 |
| *Cosmarium biretum* | GenBank: AM920339.1 |
| *Cosmarium botrytis* | GenBank: AM920378.1 |
| *Cosmarium crenatum* | GenBank: AM920370.1 |

**Table S21** – *Continued from previous page*

| Species | Accession ID |
|---|---|
| *Cosmarium meneghinii* | GenBank: AM920366.1 |
| *Cosmarium punctulatum* | GenBank: AM920373.1 |
| *Cosmarium subprotumidum* | GenBank: AM920375.1 |
| *Crocosphaera watsonii* | GenBank: AY620237.1 |
| *Cryptomonas curvata* | GenBank: JX490049.1 |
| *Cryptomonas erosa* | GenBank: AM396361.1 |
| *Cryptomonas marssonii* | GenBank: EU163586.1 |
| *Cryptomonas ovata* | GenBank: KC928318.1 |
| *Cryptomonas pyrenoidifera* | GenBank: AJ566180.1 |
| *Cucumis sativus* | GenBank: AF206894.1 |
| *Cyclotella cryptica* | GenBank: AY485499.1 |
| *Cyclotella meneghiniana* | GenBank: HM805030.1 |
| *Cylindrospermopsis raciborskii* | GenBank: AF516730.1 |
| *Cylindrotheca closterium* | GenBank: GQ468542.1 |
| *Desmidium swartzii* | GenBank: AJ428133.1 |
| *Desmodesmus abundans* | GenBank: KC861671.1 |
| *Desmodesmus maximus* | GenBank: KJ094574.1 |
| *Detonula confervacea* | GenBank: HQ912617.1 |
| *Diapensia lapponica* | GenBank: AF419794.1 |
| *Diatoma tenue* | GenBank: AJ535143.1 |
| *Dinobryon divergens* | GenBank: EU025020.1 |
| *Ditylum brightwellii* | GenBank: X85386.2 |
| *Dolichospermum flosaquae* | GenBank: AB042858.1 |
| *Dunaliella bioculata* | GenBank: DQ009761.1 |
| *Dunaliella primolecta* | GenBank: DQ009764.1 |
| *Dunaliella salina* | GenBank: DQ447648.1 |
| *Dunaliella tertiolecta* | GenBank: EF473747.1 |
| *Dunaliella viridis* | GenBank: DQ009776.1 |
| *Egeria densa* | GenBank: JF975484.1 |
| *Elodea canadensis* | GenBank: AF168841.1 |
| *Enteromorpha intestinalis* | GenBank: AJ000040.1 |
| *Eucampia zodiacus* | GenBank: KC309495.1 |
| *Euglena gracilis* | GenBank: AY029409.1 |
| *Eutreptiella gymnastica* | GenBank: FJ719618.1 |
| *Eutreptiella pomquetensis* | GenBank: AJ532398.1 |
| *Fibrocapsa japonica* | GenBank: AY788931.1 |
| *Fontinalis antipyretica* | GenBank: AF023714.1 |
| *Fragilaria barbararum* | GenBank: AJ971376.1 |
| *Fragilaria bidens* | GenBank: AM497732.1 |
| *Fragilaria capucina* | GenBank: EF465471.1 |
| *Fragilaria crotonensis* | GenBank: AM712616.1 |
| *Fragilariopsis cylindrus* | GenBank: EF140624.1 |
| *Fragilariopsis kerguelensis* | GenBank: KJ866919.1 |
| *Gambierdiscus toxicus* | GenBank: EF202890.1 |

**Table S21** – *Continued from previous page*

| Species | Accession ID |
|---|---|
| *Gephyrocapsa oceanica* | GenBank: KC404159.1 |
| *Glycine max* | GenBank: X02623.1 |
| *Gonatozygon monotaenium* | GenBank: AJ428084.1 |
| *Gonyostomum semen* | GenBank: AB512123.1 |
| *Gossypium hirsutum* | GenBank: L24145.1 |
| *Grammonema striatula* | GenBank: X77704.1 |
| *Guinardia flaccida* | GenBank: AJ535191.1 |
| *Gymnodinium breve* | GenBank: AF172714.1 |
| *Gymnodinium catenatum* | GenBank: AF022193.1 |
| *Gymnodinium mikimotoi* | GenBank: AF022195.1 |
| *Gymnodinium sanguineum* | GenBank: AJ415513.1 |
| *Gyrodinium aureolum* | GenBank: AF172713.1 |
| *Gyrodinium instriatum* | GenBank: DQ084522.1 |
| *Gyrodinium uncatenum* | GenBank: EF492498.1 |
| *Haematococcus pluvialis* | GenBank: JQ315539.1 |
| *Haptolina ericina* | GenBank: AM491030.2 |
| *Haptolina hirta* | GenBank: AJ246272.1 |
| *Helianthus annuus* | GenBank: AF107577.1 |
| *Heterocapsa triquetra* | GenBank: AF022198.1 |
| *Heterosigma akashiwo* | GenBank: AB217869.1 |
| *Hordeum vulgare* | GenBank: AY552749.1 |
| *Hydrilla verticillata* | GenBank: KM982363.1 |
| *Ipomoea batatas* | GenBank: HM053485.1 |
| *Isochrysis galbana* | GenBank: AJ246266.1 |
| *Katodinium rotundatum* | GenBank: AF274267.1 |
| *Koliella antarctica* | GenBank: AJ311569.1 |
| *Lactuca sativa* | GenBank: M82530.1 |
| *Lantana camara* | GenBank: AJ236049.1 |
| *Larix decidua* | GenBank: AB026938.1 |
| *Larrea tridentata* | GenBank: AY929372.1 |
| *Lauderia annulata* | GenBank: DQ514849.1 |
| *Lemna minor* | GenBank: S67398.1 |
| *Leptocylindrus danicus* | GenBank: AJ535175.1 |
| *Leptolyngbya tenuis* | GenBank: GQ859652.1 |
| *Limnothrix redekei* | GenBank: FM177493.1 |
| *Lingulodinium polyedrum* | GenBank: AB693195.1 |
| *Liriodendron tulipifera* | GenBank: AF206954.1 |
| *Lithophyllum margaritae* | GenBank: KP192392.1 |
| *Lobochlamys segnis* | GenBank: AJ410456.1 |
| *Lolium perenne* | GenBank: AY519271.1 |
| *Mallomonas acaroides* | GenBank: JX946333.1 |
| *Mallomonas caudata* | GenBank: U73228.1 |
| *Mallomonas crassisquama* | GenBank: KM817866.1 |
| *Mallomonas elongata* | GenBank: GU935621.1 |

**Table S21** – *Continued from previous page*

| Species | Accession ID |
|---|---|
| *Mallomonas tonsurata* | GenBank: GU935620.1 |
| *Mastigocladus laminosus* | GenBank: DQ431003.1 |
| *Merismopedia tenuissima* | GenBank: AJ639891.1 |
| *Mesotaenium kramstae* | GenBank: AJ553922.1 |
| *Micractinium pusillum* | GenBank: AM231738.1 |
| *Micrasterias americana* | GenBank: FR852595.1 |
| *Microcoleus vaginatus* | GenBank: EF654062.1 |
| *Microcystis aeruginosa* | NCBI Reference Sequence: NR_074314.1 |
| *Microcystis ichthyoblabe* | GenBank: AJ635433.1 |
| *Microcystis viridis* | GenBank: U40332.2 |
| *Microcystis wesenbergii* | GenBank: U40334.1 |
| *Micromonas pusilla* | GenBank: KP899802.1 |
| *Monodus subterranea* | GenBank: KF848930.1 |
| *Monoraphidium contortum* | GenBank: AY846375.1 |
| *Monoraphidium convolutum* | GenBank: AY846377.1 |
| *Monoraphidium griffithii* | GenBank: AY846378.1 |
| *Mucidosphaerium pulchellum* | GenBank: AY323838.1 |
| *Mucuna pruriens* | GenBank: AF525695.1 |
| *Mychonastes homosphaera* | GenBank: X73996.1 |
| *Nannochloris atomus* | GenBank: AB080303.1 |
| *Nannochloropsis oceanica* | GenBank: FJ896231.1 |
| *Nannochloropsis oculata* | GenBank: AF045045.1 |
| *Navicula arenaria* | GenBank: KJ961668.1 |
| *Navicula pelliculosa* | GenBank: AY485454.1 |
| *Nerium oleander* | GenBank: AF107572.1 |
| *Nicotiana tabacum* | GenBank: AJ236016.1 |
| *Nitzschia dissipata* | GenBank: AJ867018.1 |
| *Nitzschia frigida* | GenBank: JQ582669.1 |
| *Nitzschia paleacea* | GenBank: AJ866996.1 |
| *Nitzschia sigma* | GenBank: AJ867279.1 |
| *Nostoc muscorum* | GenBank: HF678509.1 |
| *Nothofagus grandis* | GenBank: GU476452.1 |
| *Odontella aurita* | GenBank: HQ912687.1 |
| *Odontella mobiliensis* | GenBank: KC309500.1 |
| *Odontella regia* | GenBank: KC309502.1 |
| *Odontella sinensis* | GenBank: HQ912564.1 |
| *Olea europaea* | GenBank: L49289.1 |
| *Olisthodiscus luteus* | GenBank: AY788937.1 |
| *Oryza sativa* | GenBank: AF069218.1 |
| *Ostreopsis ovata* | GenBank: AF244939.1 |
| *Pandorina morum* | GenBank: JQ315554.1 |
| *Pavlova lutheri* | GenBank: AF102369.1 |
| *Pediastrum boryanum* | GenBank: AY663036.1 |
| *Pediastrum duplex* | GenBank: M62997.1 |

**Table S21** – *Continued from previous page*

| Species | Accession ID |
|---|---|
| *Pelagomonas calceolata* | GenBank: EF455763.1 |
| *Peridinium bipes* | GenBank: DQ372891.1 |
| *Phaeocystis antarctica* | GenBank: JN381495.1 |
| *Phaeocystis globosa* | GenBank: AY851301.1 |
| *Phaeocystis pouchetii* | GenBank: AJ278036.1 |
| *Phaeodactylum tricornutum* | GenBank: GQ452861.1 |
| *Picea mariana* | GenBank: L01782.1 |
| *Picochlorum oculatum* | GenBank: AY422075.1 |
| *Pinus elliottii* | GenBank: AF051798.1 |
| *Planktothrix agardhii* | GenBank: FJ159128.1 |
| *Planktothrix mougeotii* | GenBank: FJ434250.1 |
| *Plantago lanceolata* | GenBank: AJ236046.1 |
| *Pleodorina californica* | GenBank: FJ610145.1 |
| *Pleurotaenium trabecula* | GenBank: AJ428131.1 |
| *Populus tremuloides* | GenBank: AF206999.1 |
| *Porphyridium purpureum* | GenBank: AB045584.1 |
| *Posidonia australis* | GenBank: GQ497582.1 |
| *Posidonia oceanica* | GenBank: AY491942.1 |
| *Potamogeton perfoliatus* | GenBank: AY952389.1 |
| *Proboscia indica* | GenBank: AY485470.1 |
| *Proboscia inermis* | GenBank: EF192984.1 |
| *Prochlorococcus marinus* | GenBank: AF180967.1 |
| *Prorocentrum concavum* | GenBank: Y16237.1 |
| *Prorocentrum dentatum* | GenBank: AY551273.1 |
| *Prorocentrum donghaiense* | GenBank: AJ841810.1 |
| *Prorocentrum gracile* | GenBank: AY443019.1 |
| *Prorocentrum lima* | GenBank: Y16235.1 |
| *Prorocentrum mexicanum* | GenBank: Y16232.1 |
| *Prorocentrum micans* | GenBank: AJ415519.1 |
| *Prorocentrum minimum* | GenBank: JF715165.1 |
| *Prunus persica* | GenBank: L28749.1 |
| *Prymnesium parvum* | GenBank: AJ246269.1 |
| *Prymnesium polylepis* | GenBank: AJ004866.1 |
| *Pseudochattonella farcimen* | GenBank: AM075624.1 |
| *Pseudochattonella verruculosa* | GenBank: AM075625.1 |
| *Pseudodidymocystis planctonica* | GenBank: AB037087.1 |
| *Pseudo-nitzschia granii* | GenBank: GU373962.1 |
| *Pseudo-nitzschia multiseries* | GenBank: AM235382.1 |
| *Pseudo-nitzschia pseudodelicatissima* | GenBank: GU373965.1 |
| *Pseudo-nitzschia seriata* | GenBank: GU373969.1 |
| *Pycnococcus provasolii* | GenBank: X91264.1 |
| *Pyramimonas disomata* | GenBank: FN562440.1 |
| *Pyrodinium bahamense* | GenBank: DQ500120.1 |
| *Quercus rubra* | GenBank: AF132892.1 |

**Table S21** – *Continued from previous page*

| Species | Accession ID |
|---|---|
| *Quercus suber* | GenBank: GU476438.1 |
| *Ranunculus acris* | GenBank: M82509.1 |
| *Raphidocelis subcapitata* | GenBank: HM483520.1 |
| *Rhizosolenia robusta* | GenBank: AY485481.1 |
| *Rhizosolenia setigera* | GenBank: AY485461.1 |
| *Rhodomonas salina* | GenBank: HM126532.1 |
| *Rosa hybrida* | GenBank: X66773.1 |
| *Roya anglica* | GenBank: AJ428081.1 |
| *Ruppia maritima* | GenBank: JN034103.1 |
| *Scenedesmus acuminatus* | GenBank: AB037088.1 |
| *Scenedesmus acutus* | GenBank: AJ249512.1 |
| *Scenedesmus dimorphus* | GenBank: KC790431.1 |
| *Scenedesmus quadricauda* | GenBank: KC790429.1 |
| *Scrippsiella trochoidea* | GenBank: EF492513.1 |
| *Selenastrum minutum* | GenBank: AY846380.1 |
| *Skeletonema ardens* | GenBank: DQ396522.1 |
| *Skeletonema costatum* | GenBank: JF489959.1 |
| *Skeletonema japonicum* | GenBank: DQ011160.1 |
| *Skeletonema marinoi* | GenBank: JF489953.1 |
| *Skeletonema menzelii* | GenBank: AJ535168.1 |
| *Skeletonema pseudocostatum* | GenBank: X85393.1 |
| *Skeletonema tropicum* | GenBank: EF138941.1 |
| *Solanum lycopersicum* | GenBank: KJ813722.1 |
| *Solanum tuberosum* | GenBank: FJ710157.1 |
| *Solenostemon scutellarioides* | GenBank: EU019244.1 |
| *Sorghum bicolor* | GenBank: M82328.1 |
| *Sphaerospermopsis aphanizomenoides* | GenBank: GU197654.1 |
| *Sphagnum angustifolium* | GenBank: GQ375058.1 |
| *Sphagnum squarrosum* | GenBank: GQ375075.1 |
| *Spinacia oleracea* | GenBank: L24420.1 |
| *Spirulina platensis* | GenBank: AB074508.1 |
| *Staurastrum avicula* | GenBank: EF507555.1 |
| *Staurastrum pingue* | GenBank: AJ428109.1 |
| *Staurodesmus cuspidatus* | GenBank: EF507538.1 |
| *Stellarima microtrias* | GenBank: EU090011.1 |
| *Stephanodiscus hantzschii* | GenBank: DQ093370.1 |
| *Stephanopyxis palmeriana* | GenBank: AY485527.1 |
| *Synechococcus elongatus* | GenBank: HF678511.1 |
| *Synechococcus lividus* | GenBank: AF132772.1 |
| *Synedra acus* | GenBank: AM497723.1 |
| *Synura petersenii* | GenBank: U73223.1 |
| *Synura sphagnicola* | GenBank: U73221.1 |
| *Syracosphaera pulchra* | GenBank: AM490987.2 |
| *Thalassionema nitzschioides* | GenBank: X77702.2 |

**Table S21** – *Continued from previous page*

| Species | Accession ID |
|---|---|
| *Thalassiosira allenii* | GenBank: HM991688.1 |
| *Thalassiosira curviseriata* | GenBank: AJ810859.1 |
| *Thalassiosira eccentrica* | GenBank: X85396.1 |
| *Thalassiosira guillardii* | GenBank: AF374478.2 |
| *Thalassiosira hendeyi* | GenBank: AM050629.1 |
| *Thalassiosira nordenskioeldii* | GenBank: DQ093365.1 |
| *Thalassiosira oceanica* | GenBank: DQ093364.1 |
| *Thalassiosira pseudonana* | GenBank: AY485452.1 |
| *Thalassiosira rotula* | GenBank: AF374480.2 |
| *Thalassiosira weissflogii* | GenBank: AY485445.1 |
| *Trichodesmium erythraeum* | NCBI Reference Sequence: NR_074275.1 |
| *Trichormus variabilis* | GenBank: DQ234833.1 |
| *Triticum aestivum* | GenBank: AY049040.1 |
| *Tychonema bourrellyi* | GenBank: FJ184385.1 |
| *Ulva lactuca* | GenBank: KF419328.1 |
| *Vallisneria americana* | GenBank: AF069201.1 |
| *Vitis vinifera* | GenBank: GQ849399.1 |
| *Zea mays* | GenBank: AF168884.1 |
| *Zostera marina* | GenBank: HQ445940.1 |
| *Zostera noltii* | GenBank: AF207058.1 |
